# Functional analyses and integrated mechanisms of cellular destruction by L-amino acid oxidase

**DOI:** 10.1101/2024.09.16.613219

**Authors:** Krisna Prak, Christin Luft, Eliona Tsefou, Carlos Chávez-Olórtegui, Janos Kriston-Vizi, Robin Kettler, Vania M. M. Braga

## Abstract

Snakebite accidents are prevalent worldwide and cause a spectrum of severe clinical manifestations and reduction of patient quality of life and economic income. L-amino acid oxidase (LAAO) is a highly toxic enzyme present in various venoms that causes tissue necrosis, oedema, coagulopathies, and organ failure. Here we investigate the mechanisms of LAAO cytotoxicity preceding cell death using recombinant LAAO and a catalytic inactive LAAO mutant. Wild-type LAAO uptake by cells leads to a decrease in lysosome number and size and inhibition of autophagy flux. Mitochondria function is impaired by severe proton leakage and mitochondrial fission is stimulated. Despite engulfment by autophagosomes, mitochondrial clearance is prevented by the lysosomal defects. The coordinate multi-organelle dysfunction strongly perturbs energy production, cell metabolism and clearance of defective organelles by autophagy, thereby triggering an irreversible destructive path. Considering the fast organelle impairment, strategies to reduce multi-organelle injury after LAAO exposure may be effective to maintain critical cell functions and strengthen adaptive responses against cytotoxicity.

## INTRODUCTION

Compared to other neglected tropical diseases, snakebites have similar global prevalence to cholera infection ^1^ and yet show the largest patient lethality, disproportionally affecting the poorest communities in developing countries ^2^. The highest impact of snakebite accidents is seen on the severe, debilitating disabilities that often cause a life-long reduction of income and quality of life with extensive necrosis, disfigurement, and amputations ^1,3^. Available treatment for envenoming patients relies on quenching of circulating venom toxins with antivenom sera (when available in hospitals and with appropriate snake species specificity) and the natural elimination of toxins from the body. Because of the high costs and low availability of anti-venoms worldwide, there is currently an unmitigated clinical need to develop novel treatments for snakebite envenoming ^4,5^, particularly the devastating necrosis at the snakebite site where anti-venoms are not effective.

A concerted effort in the field has been to characterize the structure and biochemistry of the clinically relevant venom toxins. Yet, the lack of detailed cellular mechanisms of how specific toxins promote cell death is a major hurdle in the field. Understanding the cellular action of venom toxins will provide insights to regenerate tissue damage and accelerate patient recovery. Whether the specific responses elicited are a direct consequence of the action of venom toxins or a defensive mechanism to protect against cytotoxicity is unclear. It is indeed feasible that cell adaptive responses against toxicity may compound the severe reduction in cell fitness and accelerate cell death ^6^.

To address the above conceptual gap, we performed a structure-function analyses of snake venom L-amino acid oxidase (LAAO, E.C.1.4.3.2) to inform its cellular mechanisms. Purified native venom LAAO causes local tissue damage, inhibition and induction of platelet aggregation, haemorrhage, haemolysis, and oedema in animal models ^7^. LAAO proteins are found from bacteria to mammals. They have conserved catalytic pockets and are thought to participate in amino acid catabolism and defensive mechanisms ^8^. Mammalian LAAOs are found in the brain, milk, sperm and immune cells. Among mammalian LAAOs, Interleukin-4 induced gene 1 (IL4I1) is the best characterized protein; it is secreted at human immune synapses ^9^ and contributes to immunoregulatory responses. While both human LAAO and snake venom LAAO oxidize L-amino acids, only the venom counterpart is cytotoxic ^10^.

Snake venom LAAOs are flavoproteins that have preferential catalytic specificity for hydrophobic and aromatic L-amino acids ^7,11^. After oxidation of L-amino acid substrate, the catalysis by-products (ammonia and hydrogen peroxide are toxic *per se*. Hydrogen peroxide can be converted to reactive oxygen species (ROS), i.e., highly reactive hydroxyl radical or intracellular superoxide that have major effects in various cellular processes ^12^. Whether LAAO cytotoxicity is triggered by alterations in amino acid catabolism and exacerbated by the oxidative stress responses is an attractive possibility that is currently underexplored. For example, oxidative stress is a key feature of envenomation by both purified toxins and crude venom from various snake species ^13,14^.

The specific cellular mechanisms underlying snake venom LAAO toxicity are poorly understood. We found that purified native LAAO is internalized by primary human cells and causes cytoplasmic vesicles swelling, pyknotic nuclei, cell retraction, induction of autophagy followed by apoptosis and necrosis ^15^. Autophagy is a cellular process that regulates catabolism, homeostasis and quality control of organelles ^16^. During intoxication by different snake venoms or purified toxins ^15,17^, autophagy may be a common denominator of cellular envenomation.

LAAO toxicity ^15^ likely alters cellular homeostasis processes essential for cell survival. We hypothesis that LAAO catalysis interferes directly or indirectly with the functions of mitochondria and lysosomes, two essential organelles. Mitochondria participate in programmed cell death regulation and have a key function in energy production, ROS generation and ROS metabolism ^18,19^. Mitochondrial quality control is essential for the maintenance of cellular homeostasis: damaged mitochondria is removed via mitophagy to maintain cell fitness ^20^. Dysfunctional lysosomes are associated with various pathological status ^21^ as lysosomes are key regulators of degradative processes, cell secretion, metabolic sensing and adaptation ^22^.

We envisage three possibilities. First, the catalyses by-product ammonia may inhibit lysosomal fusion with other organelles ^23^, while oxidative stress generated by its catalysis could damage mitochondria. Second, LAAO is expected to oxidise and reduce the levels of its amino acid substrates, which may alter cell metabolic rate. Third, it is also feasible that the unbalancing amino acid levels may affect lysosomal function, as lysosomes form a platform for activation of the amino acid sensing machinery ^24^.

To overcome the restricted availability of purified native toxins from snake venoms ^15^, we produced active recombinant LAAO from *Bothrops atrox* (*B. atrox*) snake, the species responsible for most accidents in the Brazilian Amazon ^3,25^. Available crystal structures of native LAAO from different snake species detail a remarkable domain conservation of a helical domain that provides access to the active site and the binding domains of the co-factor flavin-adenine dinucleotide (FAD) and substrate ^26–28^. Yet, conserved residues that participate in LAAO catalysis ^27^ have not yet been formally identified (apart from one study^10^).

Using various LAAO mutants and kinetic analyses, we demonstrate unequivocally the key amino acid residues important for catalysis and confirm their contribution *in vitro* and *in cellulo*. A comprehensive temporal analyses of LAAO-induced cellular phenotypes show that LAAO induces cell death by targeting mitochondrial and lysosomal function and dynamics, and impairs catabolic processes, energy production and consumption. Our data shed insights into the LAAO-dependent early cytotoxic events and provide a platform to investigate the factors underpinning the rapid cell death upon exposure to snake toxins. Our data will be instrumental to increase our conceptual knowledge of toxins on their own right and to identify strategies to ameliorate cell destruction and preserve tissue integrity.

## RESULTS

### Insights into the structure of LAAO catalytic site

Snake venom LAAOs have very similar amino acid sequence (Fig. S1). *B. atrox* venom LAAO monomeric structure (Fig. 1A) is most similar to that of *Agkistrodon halys pallas* (*A. H. pallas*) venom LAAO (RMSD 0.38 Å using PyMOL analysis). We selected a few amino acid residues to investigate among several amino acid residues proposed to form LAAO catalytic pocket or have a close interaction with substrate and cofactor FAD (Fig. 1B). Arginine residue (R90) has been suggested to interact with the co-factor flavin-adenine dinucleotide (FAD) and the substrate ^28^, while tyrosine at position 372 (Y372) is proposed as a substrate/ligand binding residue ^29^. A glycosylated site at an asparagine residue (N172) seems important for catalysis (Fig. 1A), albeit a surface residue ^28^. Finally, R322 and H223 have been proposed as important residues in the catalytic pocket ^29^. We produced various recombinant proteins mutated in these residues and analysed their catalysis efficiency in the oxidation reaction (Fig. 1C) ^7^ and *in cellulo* effects (see below).

**Fig. 1.**
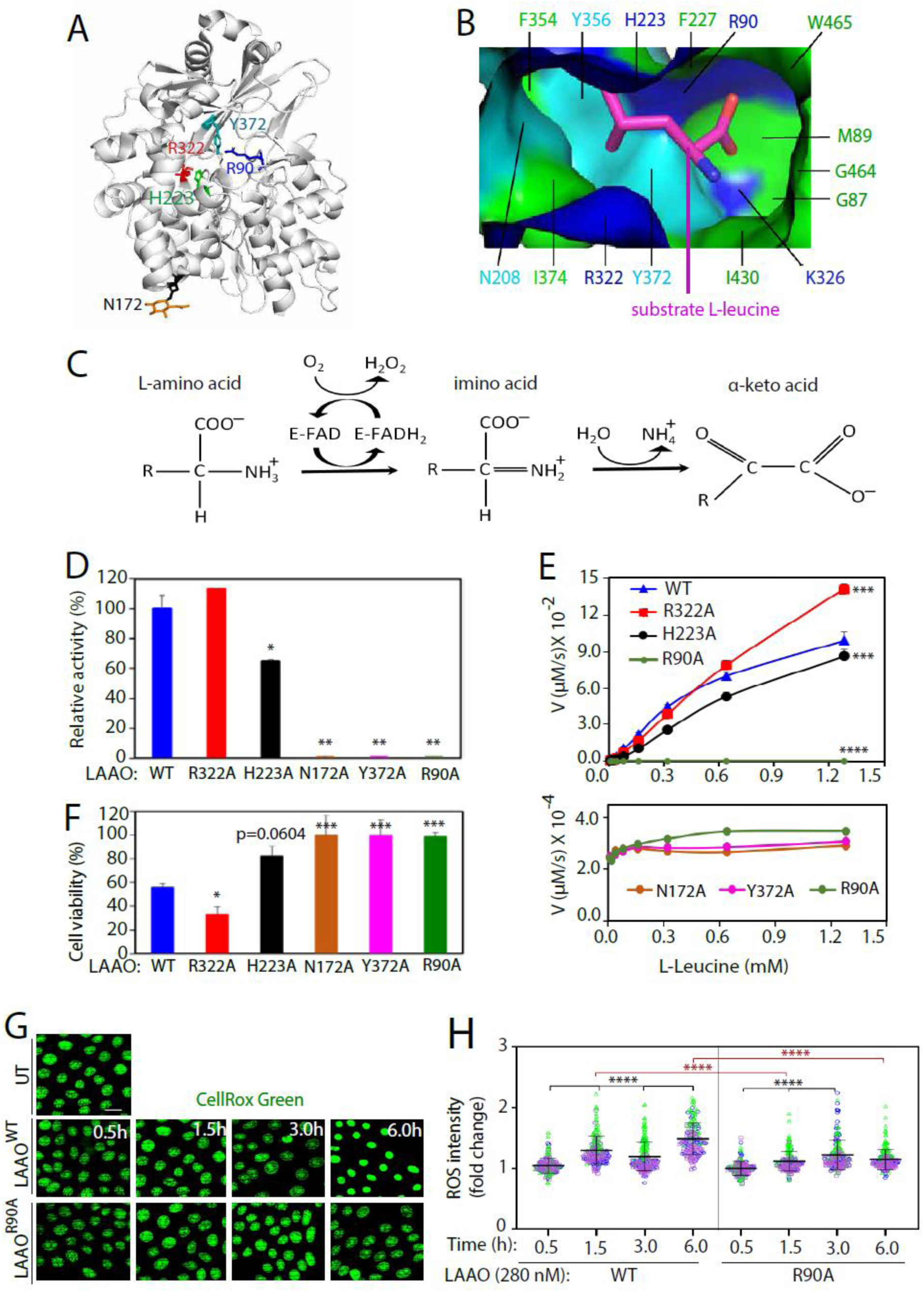
Catalytic activity, cell cytotoxicity, and oxidative stress of recombinant *B. atrox* LAAO^WT^ and mutants. **A**, Monomeric structure of *B. atrox* LAAO showing the predicted catalytic and substrate binding residues as sticks: Y372, R322, R90, H223, N172. **B**, Surface of the substrate binding pocket of LAAO (*A. H. pallas* venom (PDB 1TDN) showing positively charged amino acid residues (blue), polar residues (cyan), and hydrophobic residues (green). L-leucine substrate is shown as magenta stick. **C**, L-amino acid oxidation reaction catalysed by LAAO^WT^ with co-factor FAD (E-FAD) generating H_2_O_2_ and ammonia as by products. **D**-**E,** Activity of purified LAAO^WT^ or LAAO mutants (52.72 nM) compared to wild type (D, arbitrarily set at 100%) or their initial velocity V (E; µM per second, Y axis). The initial velocity V is defined as the change in the concentration of H_2_O_2_ produced by LAAO at each concentration of substrate L-leucine (mM, X axis). In the bottom panel, catalytic activities of selected mutants are shown magnified (100X). **F**, Keratinocyte cell viability was assessed by incubation 14 nM of LAAO^WT^ or mutants. **G-H**, Cellular reactive oxygen species (ROS) generated by keratinocytes untreated or treated with 280 nM LAAO^WT^ or LAAO^R90A^ and live stained with CellRox Green (green) and fixed before confocal imaging. Representative images (G) and quantification (H) are shown. Bars indicate mean and error bars show standard deviation. The assays were done as multiple technical replicates in three independent biological replicates (thereafter N=3), represented in blue, green, or pink colours. Black asterisks represent comparison within a single treatment group; red asterisk compares across different treatments. **** P ≤ 0.0001, *** P ≤ 0.001, ** P ≤ 0.01, * P ≤ 0.05. Number of all images used for the analysis are indicated in File S1. Scale bar: 20 µm.

### *B. atrox* LAAO catalytic kinetics reveals the essential role of residues R90, Y372 and N172

The production of recombinant active LAAO has its challenges due to its high toxicity, insolubility in bacteria, distinct glycosylation by yeast ^27,30,31^ or limited secretion in mammalian systems ^10,27^. Using HEK293T cells, we successfully produced recombinant LAAO and mutated versions of the protein on the proposed residues for LAAO catalyses (R322, H223, N172, Y372 and R90; Fig. S2A).

Among the preferred *in vitro* substrates for native *B. atrox* LAAO (L-Tyr, L-Phe, L-Ala, and L-Leu) ^11^, we chose L-Leu to determine the *in vitro* catalytic activity of LAAO recombinant proteins. The mutants LAAO^R90A^, LAAO^N172A^ and LAAO^Y372A^ showed a strong reduction in their activity (Fig. 1D), while LAAO^H223A^ showed partial inhibition of catalysis (Fig. 1D) as previously reported ^31^. Unexpectedly, LAAO^R322A^ had a small, but reproducible activity when compared to LAAO^WT^. Catalytic kinetics (Fig 1E) showed that the enzymatic velocity of LAAO^R322A^ (at 26.36 nM) is slower than the LAAO^WT^ (L-Leu concentration at ≤ 300 µM) but became faster at substrate concentrations above 600 µM. The catalytic efficiency (K_cat_/ K_m_) of recombinant LAAO^WT^ and the relative catalytic activity was strongly reduced for all mutants, apart from LAAO^R322A^ (Table 1). The contribution of residue R322 to catalysis remains unclear, and it will be investigated elsewhere. It is feasible that R322 may cooperate with other residues to influence catalyses, as shown with equivalent mutations in recombinant LAAO from *Naja naja* snake ^10^. Our data on *B. atrox* LAAO catalyses strengthen the case for the essential roles of R90, Y372 and N172 sites, but not R322.

**Table 1.** Enzymatic kinetics of LAAO^WT^ and its mutants. The results were derived from Figure 1E and curves fitting using the nonlinear regression method in R software. *, LAAO^R322A^ catalytic activity depends on the substrate concentration as shown in the bottom rows. Results are displayed as mean ± standard deviation. ND, not detected. N=3.

| Catalytic kinetics |  |  |  |  |  |  |
| --- | --- | --- | --- | --- | --- | --- |
| LAAO | WT | R322A | H223A | R90A | N172A | Y372A |
| Kcat (1/s) | 4.53 ± 0.32 | 17.4 ± 0.91 | 7.28 ± 1.62 | ND | ND | ND |
| Km (μM) | 501 ± 46 | 2410 ± 142 | 1457 ± 392 | ND | ND | ND |
| Kcat/Km (1/μM.s) | 9.08E-03 | * | 5.50E-03 | ND | ND | ND |
| LAAO relative activity- high L-Leucine concentration |  |  |  |  |  |  |
| LAAO | WT | R322A | H223A | R90A | N172A | Y372A |
| Activity | 1 | 1.13 ± 0.005 | 0.64 ± 0.011 | 0.0044 | 0.0041 | 0.0028 |

Consistent with native LAAO ^15^, recombinant LAAO^WT^ was toxic to normal keratinocytes (Effective Concentration (EC_50_) of 0.8 µg/ml or 14 nM; Fig. S2B). Viability assays using LAAO mutants at the same concentration (14 nM; Fig. 1F) demonstrated that LAAO^R322A^ potently reduced viability to a greater extent than LAAO^WT^, while a small reduction in cell viability was promoted by LAAO^H223A^ (Fig. 1F). The mutants N172A, Y372A or R90A did not interfere the metabolic fitness of normal keratinocytes.

We investigated the earlier LAAO cytotoxicity mechanisms leading to apoptosis and necrosis ^15^. Hydrogen peroxide is a very toxic by-product of LAAO oxidation (Fig.1C) and can be converted to reactive oxygen species (ROS). We found that in cell culture media LAAO^WT^ (28 nM) exponentially produce H_2_O_2_ to 10 mM in 6 h (Fig. S2C). To detect ROS after LAAO treatment (2xEC_50_), cells were stained with CellRox Green, a cell permeable dye (Fig. 1G-H). ROS levels fluctuated but were highest after 6 hours incubation with LAAO^WT^, while a small, transient burst of ROS levels was seen at 3 hours after LAAO^R90A^ incubation (timing and intensity distinct from LAAO^WT^). Thus, similar to native LAAO ^32^, recombinant *B. atrox* LAAO elevates cellular oxidative stress in a catalysis-dependent manner.

### LAAO disrupts autophagy flux and lysosome function

Autophagy is the earliest detectable cell stress response caused by LAAO treatment of primary cells ^15^. To investigate LAAO-induced autophagy flux, we expressed EGFP-LC3 in keratinocytes. The number of LC3 puncta (i.e., as a proxy for autophagosome numbers) at any given time is a balance between autophagosome *de novo* formation and degradation. When compared to untreated controls, LAAO^WT^ incubation significantly increased the number of LC3 puncta after 3 and 6 hours (Fig. 2A-B). In contrast, the catalytically dead mutant (LAAO^R90A^) triggered a transient surge in the number of LC3 puncta at 1.5 hours, suggesting that autophagosomes were induced and quickly degraded at later time points, returning to basal levels (Fig. 2A-B). As the number of LC3 puncta remains high after LAAO^WT^ treatment, we surmise that autophagy flux is promoted, but autophagosome degradation in lysosomes might be impaired.

**Fig. 2.**
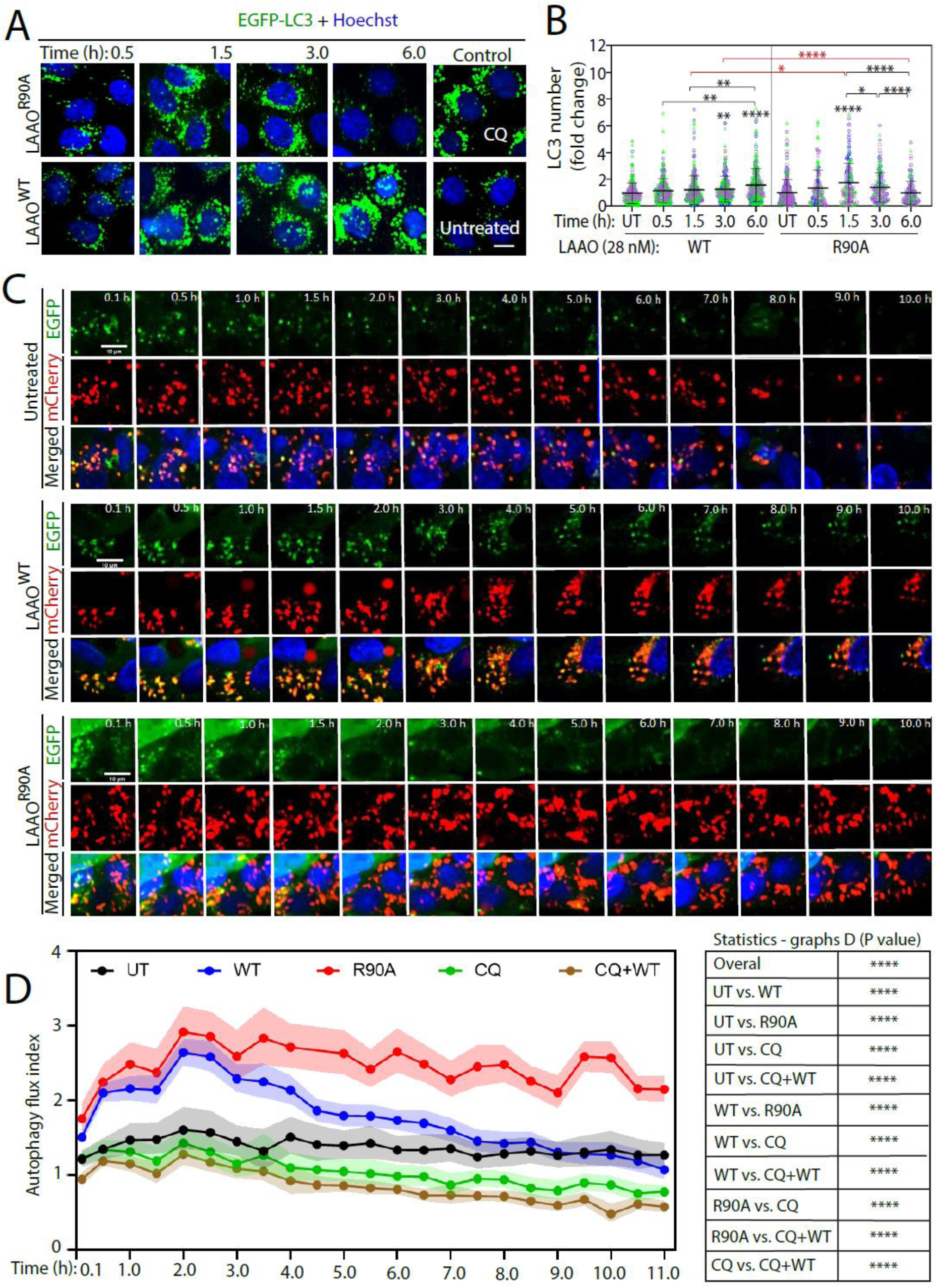
Autophagic response of keratinocytes and HepG2 cells in the presence of LAAO^WT^ or LAAO^R90A^. **A-B**, Accumulation of LC3 puncta in keratinocytes following incubation with 28 nM of either LAAO^WT^ or LAAO^R90A^ for 6 h. Controls are untreated cells or those treated with 10 μM chloroquine (CQ). Representative confocal images (A) and quantification (B) of the number of LC3 puncta in each sample is shown as normalized to untreated cells. Bars indicate mean and error bars show standard deviation. **C**, Representative confocal live imaging of HepG2 cells stable expressing tandem mCherry-EGFP-LC3 left untreated or incubated with either LAAO^WT^ or LAAO^R90A^ (1 X EC_50_, 42 nM for HepG2 cells) and images captured every 30 min. For each sample, images show the fluorescence of EGFP-LC3 (top row) or mCherry-LC3 (middle row) and the merged image (bottom row) with nuclei stained with Hoechst 33342 (blue). Representative images of controls are shown in Fig. S3: chloroquine (CQ) or chloroquine + LAAO^WT^ (CQ + WT). **D**, Autophagy flux index of HepG2. Autophagy flux index is the ratio between the number of red LC3 puncta and the number of green LC3 puncta. Graph shows geometric mean with error, 95% confidence interval (95% CI) added as light-coloured shadow (N=3). Table on the right shows the overall comparison statistics. In B, three independent biological replicates (N=3) are represented in different colours (blue, green, or pink). Multiple statistical comparisons among different groups were performed using ad hoc test (Dunn’s Method) after Kruskal–Wallis one-way ANOVA (B) or Two-way ANOVA (Tukey’s multiple comparisons test) (D). Black asterisks show comparison within a single treatment group; red asterisks show comparison across different treatments. **** P ≤ 0.0001, *** P ≤ 0.001, ** P ≤ 0.01, * P ≤ 0.05. Scale bar: 10 µm.

To differentiate between autophagosome initiation and degradation, autophagy flux analysis was performed using a different, well-established model, HepG2 cells expressing mCherry-EGFP-LC3 (Fig.2D) ^33^. Autophagy flux index is calculated as the ratio (mCherry/GFP) in LC3 puncta. It measures the rate in which autophagosomes reach the lysosomal compartment, via quenching of EGFP fluorescence (sensitive to low pH in lysosomes) while mCherry fluorescence is not affected ^33^. Basal autophagy flux index of untreated cells showed the disappearance of EGFP fluorescence in LC3 puncta as autophagosomes fuse with acidic compartment of lysosomes (Fig. 2C). Treatment with chloroquine to impair lysosome acidification and content degradation ^34^ reduced autophagy flux index throughout the time course (Fig. 2D, Fig. S3A).

Autophagy flux index was increased in LAAO^WT^- and LAAO^R90A^-treated for up to 2 hours (EC_50_ = 42 nM; Fig. 2D) ^35^, indicating higher degradation of autophagosomes in either treatment. However, after 3 hours, the autophagy flux index in LAAO^WT^ samples decreased, as clearance of autophagosomes in lysosomes was reduced, returning to basal levels with time (Fig. 2C-D). In contrast, inactive LAAO^R90A^ treatment induces a higher autophagy flux index and is significantly distinct from untreated cells (Fig.2C-D). The data suggest that the inactive LAAO mutant continues to generate autophagosome formation and effective degradation in HepG2 cells (Fig. 2D). Such continuous induction of autophagosomes by inactive LAAO treatment may reflect a catalysis-independent phenotype of clearing an extraneous exogenous protein. Finally, the higher autophagy flux caused by LAAO^WT^ treatment is blocked by chloroquine treatment (Fig. 2D, Fig. S3B), confirming that observed increase in autophagy flux index up to 3 hours requires a functional lysosome. We concluded that the catalytic activity of LAAO inhibits lysosome function and impairs autophagosome degradation.

Thus, LAAO^R90A^ phenotype of sustained high autophagy flux index in HepG2 cells (Fig. 2C-D) is consistent with the keratinocyte data (Fig. 2B) showing a transient increase of LC3 puncta likely from a fast clearance in lysosomes. Whereas the interpretation of inhibition of autophagosome clearance by LAAO^WT^ is consistent across HepG2 and keratinocyte models, it is possible that cell-specific differences may be present, due to distinct survival abilities of immortalized cell lines (HepG2) versus normal cells (keratinocytes). Overall, these set of results suggest that LAAO activity may inhibit autophagy flux either by preventing lysosomal acidification or fusion of autophagosomes with lysosomes.

Next, we investigated the intracellular localization of LAAO ^15^. In keratinocytes, recombinant His-tagged LAAO^WT^ was internalised and found in vesicles, where it partially co-localised with autophagosomes (pEGFP-LC3) and lysosomes (Lysotracker red DND-99; Fig. 3A). A progressive reduction of lysosome number and size was induced by treatment with wild-type, but not the catalytic dead LAAO (Fig. 3B-D). Taken together, we demonstrate that recombinant LAAO accumulates in autophagosomes and alters lysosome number and morphology in a catalysis-dependent manner. We surmise that the lysosome dysfunction triggered by LAAO would delay or inhibit autophagosome degradation and autophagy flux, therefore exacerbating cytotoxicity.

**Fig. 3.**
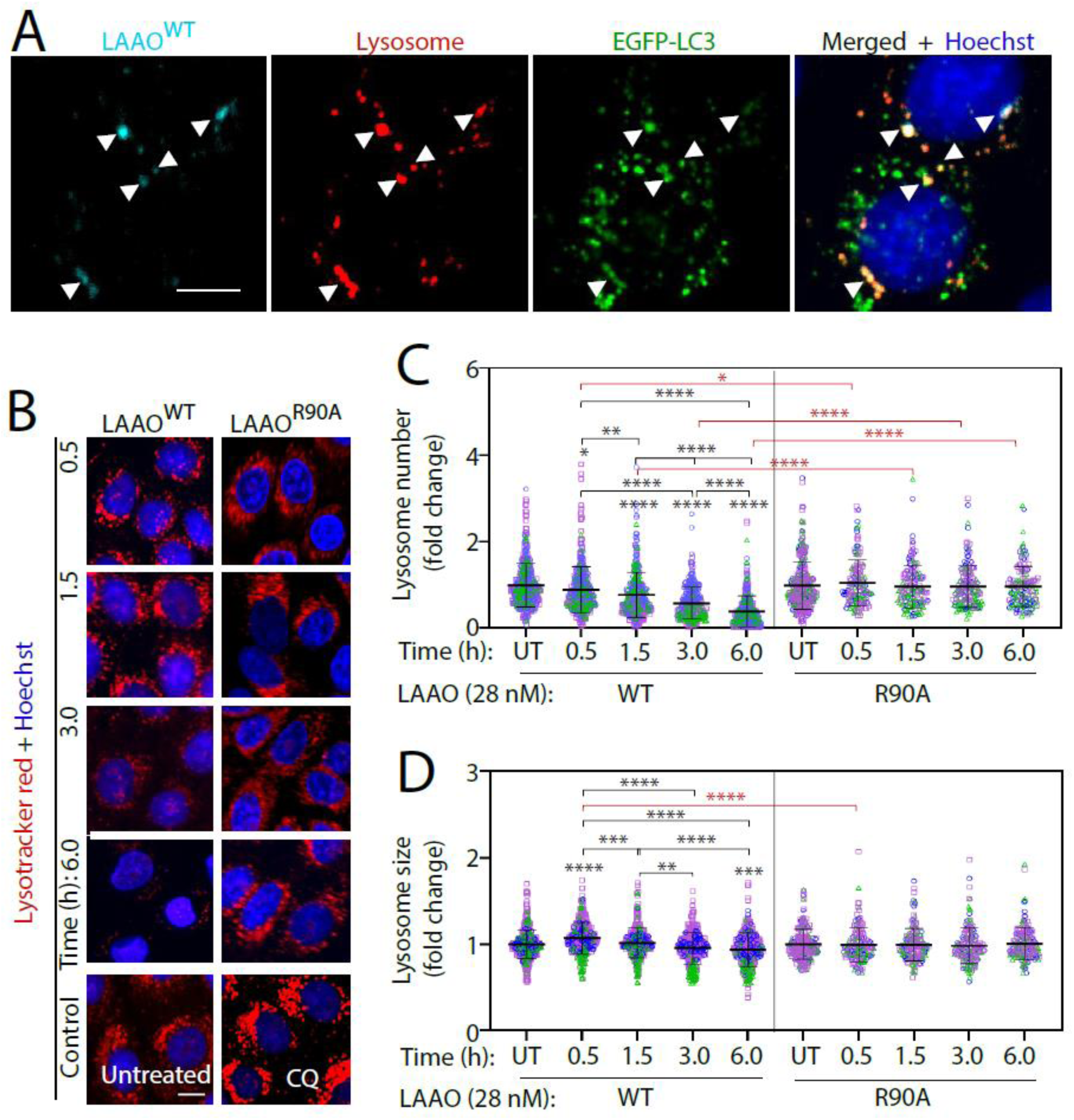
*B. atrox* LAAO internalization and lysosomal responses. **A**, Confocal images of keratinocytes transiently transfected with pEGFP-LC3 and treated with 28 nM LAAO^WT^ for 3 h. White arrowheads indicate co-localization of LAAO^WT^ (cyan) with LC3 (green) and lysosome (red). **B**-**D**, Assessment of lysosomal morphology. Keratinocytes were incubated with 28 nM of LAAO^WT^ or catalytic dead LAAO^R90A^ or 10 µM chloroquine (CQ) for up to 6 hours. Lysosomes were visualized by live staining (100 nM Lysotracker red DND-99 and 1:10000 Hoechst) for 1.5 h before the end of the treatment. Confocal live images (B) were quantified to obtain lysosome numbers (C) and sizes (D). Values are expressed as fold change relative to untreated samples. Bars indicate mean and error bars show standard deviation. N=3; independent biological replicates are represented in different colours (blue, green, or pink). Black asterisks compare within a single treatment group; red asterisks compare across different treatments. **** P ≤ 0.0001, *** P ≤ 0.001, ** P ≤ 0.01, * P ≤ 0.05. Scale bar: 10 µm.

### LAAO promotes mitochondria dysfunction and morphological changes

As the amount of LAAO in venoms from different snake species can vary extensively ^36^, higher concentrations of LAAO^WT^ (280 nM) was tested in parallel to reflect variability of LAAO concentration in different venoms. The early alterations on autophagy flux and lysosomal dynamics caused by LAAO^WT^ were not accompanied by extensive cell detachment as quantified by the nuclei number as a proxy (Fig. 4A). A significant and time-dependent decrease in cell number was most evident at 6 hours (Fig. 4A-B). Higher LAAO concentration promoted cell removal after 6 hours incubation, while the catalytically dead LAAO^R90A^ did not (Fig. 4B). Altogether, our results indicate that cell detachment induced by wild-type LAAO does not temporally correlate with the changes in autophagy flux and lysosomal function breakdown observed at earlier time points.

**Fig. 4.**
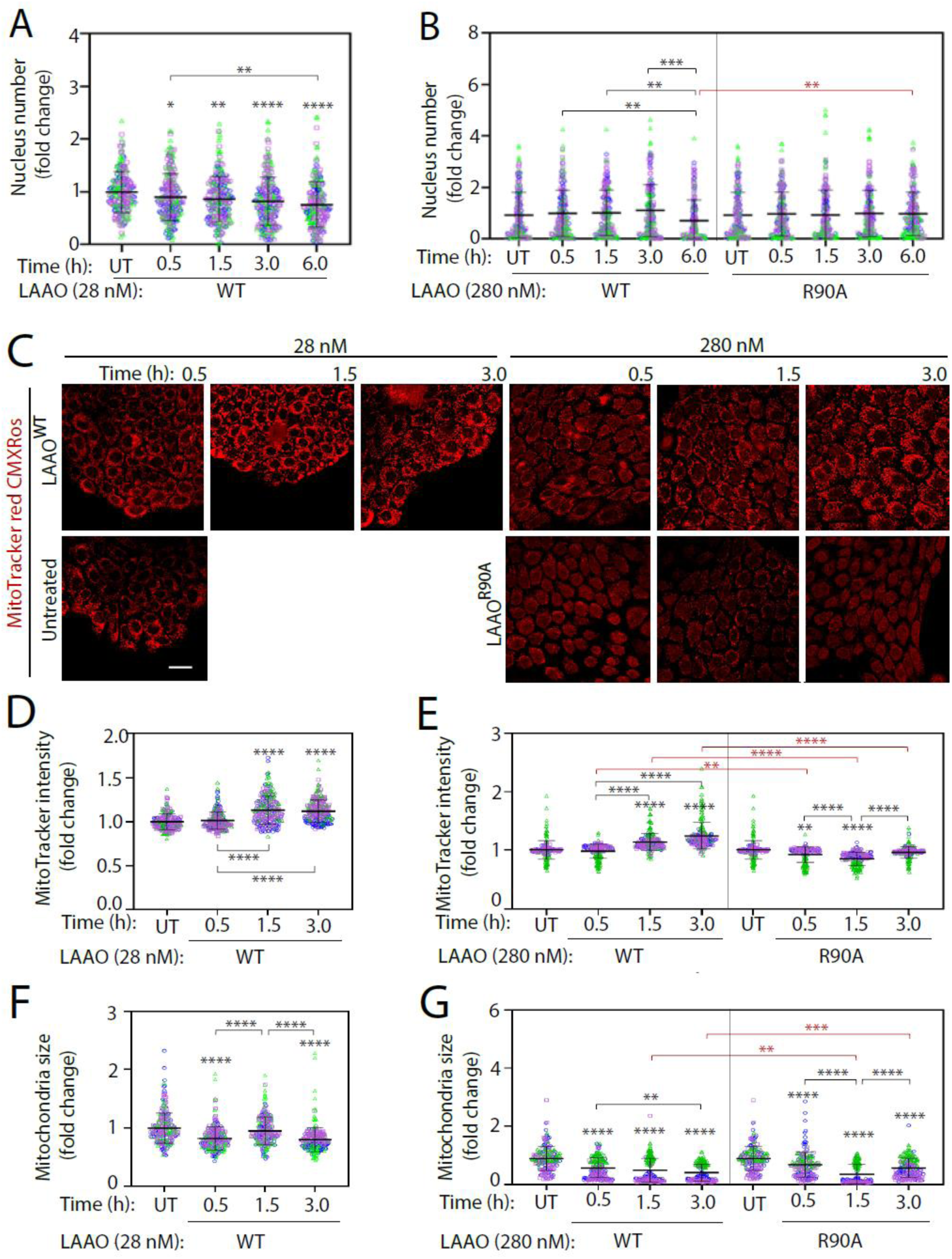
LAAO treatment reduces cell number and mitochondria size and function. Normal keratinocytes were treated with recombinant LAAO proteins at different concentrations (28 nM or 280 nM) for up to 6 hours. **A-B**, Cells were stained with 1:10000 Hoechst and the number of nuclei in each well were quantified. **C**, Cells were stained live with MitoTracker red CMXRos and fixed before confocal image acquisition. **D-G**, Images were quantified to assess mitochondria membrane potential (D-E) or mitochondria size (F–G). All graphs show values normalized to the untreated samples. Bars indicate mean and error bars show standard deviation. N=3; independent biological replicates and are represented in blue, green, or pink colours. Black asterisks compare within a single treatment group; red asterisks compare across different treatments. **** P ≤ 0.0001, *** P ≤ 0.001, ** P ≤ 0.01. Scale bar: 20 µm.

Using MitoTracker Red CMXRos (Fig. 4C-G), we found that mitochondria function was perturbed in two different ways. First, mitochondrial membrane potential was altered after treatment with LAAO^WT^ (1.5 hour onwards; Fig. 4C-E). A small, but significant increase in MitoTracker Red intensity was observed upon incubation with LAAO^WT^ (Fig. 4D-E), but not LAAO^R90A^ (Fig. 4E). Second, the membrane potential disruption by LAAO^WT^ was accompanied by a strong reduction in mitochondria size (Fig. 4F-G). At higher concentrations, catalytic inactive LAAO^R90A^ also showed similar phenotype at the first 1.5 h and had smaller mitochondria size when compared to wild-type LAAO (Fig. 4G). At 3 h, mitochondria size continued to decrease when treated with LAAO^WT^ but not with LAAO^R90A^.

Consistent with the above data, mitochondria Feret’s length (Fig. 5A-C) and mitochondria fragment number (Fig. 5D-E) were proportional to mitochondria size (Fig. 4F-G) (bigger mitochondria size ̶ longer ferret length ̶ smaller mitochondria fragment number or smaller mitochondria size ̶ shorter ferret length ̶ larger mitochondria fragment number) except the treatment at higher concentration with LAAO^WT^ at 3 h and LAAO^R90A^ at 1.5 h which resulted in smaller mitochondria size ̶ shorter ferret length ̶ smaller mitochondria fragment number, suggesting mitochondria lost. The increase in mitochondria size ̶ Feret’s length ̶ mitochondria fragment number at 3 h from 1.5 h when treated with LAAO^R90A^ indicated mitochondria recovery when treated with inactive LAAO. The reasons why inactive LAAO altered mitochondria morphology transiently in the absence of higher oxidative stress are currently unclear.

**Fig. 5.**
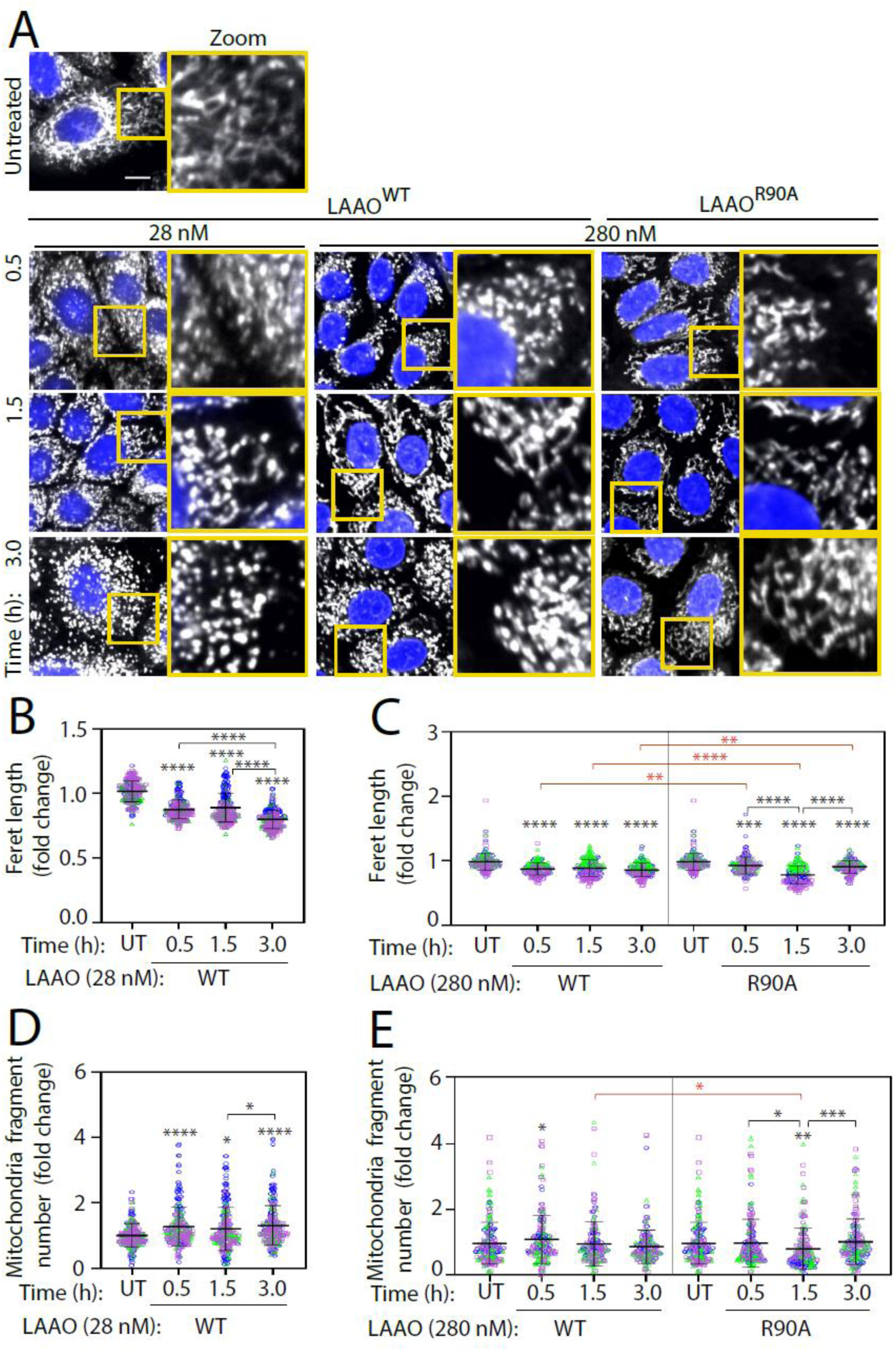
*B. atrox* LAAO alters mitochondrial network dynamics. Normal keratinocytes were treated with recombinant LAAO proteins at different concentrations (28 nM or 280 nM) for up to 3 hours. Cells were stained live with Hoeschst and MitoTracker and fixed before imaging using confocal microscopy. Images were processed to quantify mitochondria morphology (B-C) and fragmentation (D-E). All values are normalised to the untreated (UT) samples. **A**, Representative images of cells stained with MitoTracker Red CMXRos. Mitochondria and nuclei are shown in grey and blue, respectively. Yellow box shows region-of-interest that is enlarged on the right-side panel. **B**-**C**, Quantification of mitochondria Feret’s diameter (i.e., the longest distance (μm) between any two points within a given mitochondrion). **D**-**E**, Number of mitochondria fragments. Bars indicate mean and error bars show standard deviation. N=3; independent biological replicates are represented in blue, green, or pink colours. Black asterisks compare within a single treatment group; red asterisks compare across different treatments. **** P ≤ 0.0001, *** P ≤ 0.001, ** P ≤ 0.01, * P ≤ 0.05. Scale bar: 10 µm.

The discrepancy between the timing of mitochondrial fragmentation with different LAAO concentrations may be due to the ability of higher dosage of LAAO^WT^ to fragment mitochondria more potently. The phenotype with 280 nM LAAO^WT^ could result from (i) a faster and more efficient fragmentation which tails off to steady state by 1.5 hours and (ii) smaller mitochondria fragments are more readily engulfed by autophagosomes and cleared^37^. The net result was no significant increase in fragment numbers. We surmise that the high oxidative stress induced by LAAO catalysis increases mitochondria fragmentation and reduces their size and Feret’s diameter.

### Cell metabolism and energy production are strongly disturbed by LAAO

The above morphometric changes suggest that mitochondrial function is severely impaired by LAAO. As mitochondria have an essential role in generation cellular energy, we carried out bioenergetic measurements (see methods and figure legends). Following a time course, the profiles for Oxygen Consumption Rate (OCR) and Extracellular Acidification Rate (ECAR) were recorded in the presence or absence of recombinant LAAO proteins (Fig. S4). Normal keratinocytes have a preference for glycolysis ^38^, but switched to oxidative phosphorylation in glucose- and pyruvate-deprived conditions (Fig. 6A-B). LAAO^WT^ treatment impaired the ability to adapt energetically in the absence of glucose or pyruvate, while LAAO^R90A^ had no effect (Fig. 6A-B).

**Fig. 6.**
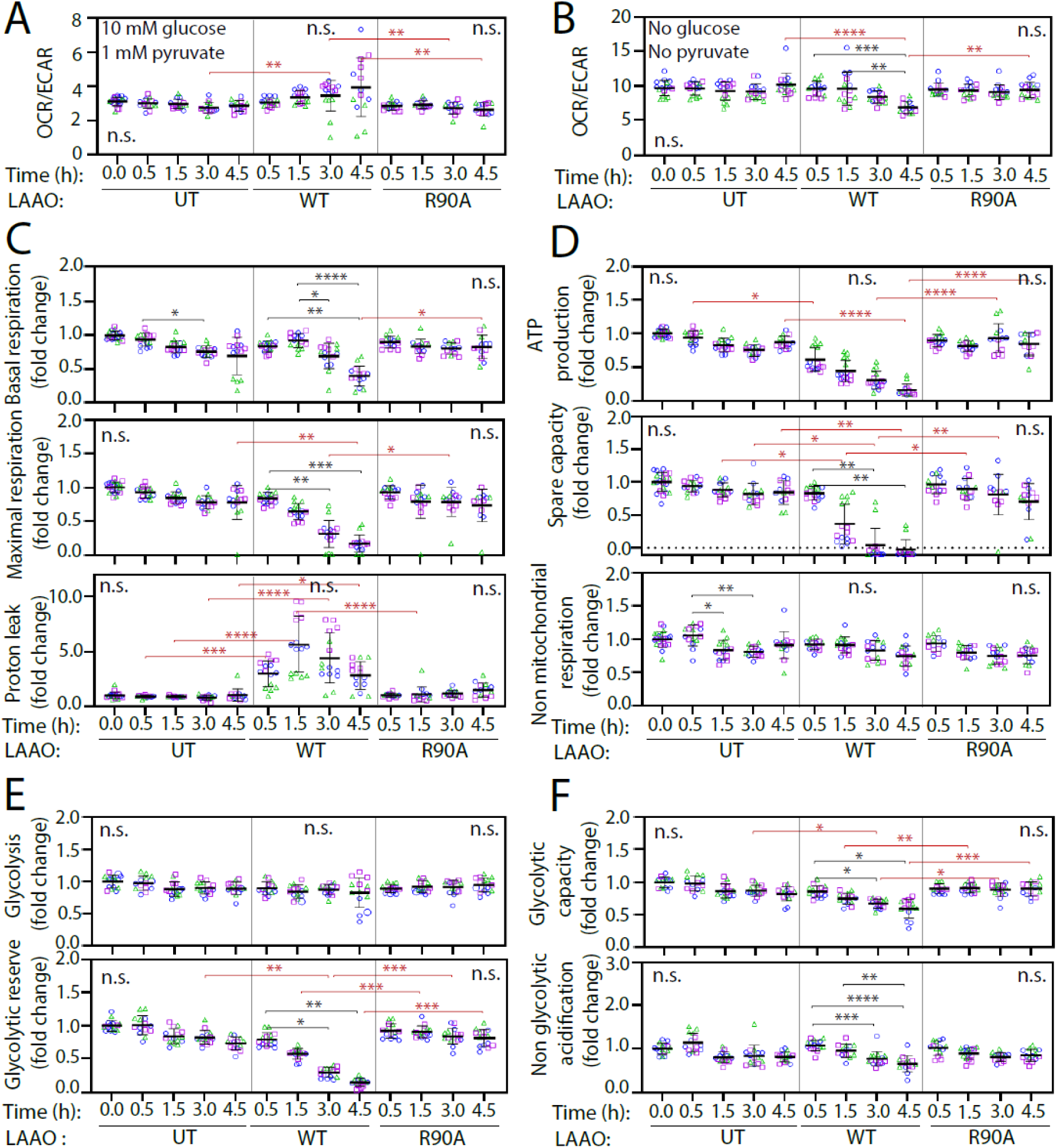
Mitochondrial dysfunction and metabolic stress are promoted by LAAO catalysis. Oxygen consumption rate (OCR) and extracellular acidification rate (ECAR) were measured in keratinocytes untreated (UT) or treated with 2X EC_50_ (73 nM for SeaHorse experiments) of LAAO^WT^ or LAAO^R90A^ for up to 4.5 h as described in Methods. Recorded OCR and ECAR profiles for Cell Mito Stress and Glycolysis Stress tests are shown in Fig. S4. **A**-**B**, Bioenergetic changes are obtained by calculating OCR/ECAR ratio at basal respiration in the presence (A) or absence (B) of glucose and pyruvate. **C**-**D**, Metabolic parameters are calculated from the Mitochondrial Stress test: non-mitochondrial and mitochondrial respiration (basal and maximal), proton leak, ATP production and spare capacity. **E**-**F**, Glycolytic parameters were calculated from the Glycolysis Stress test: glycolysis, non-glycolytic acidification, glycolytic reserve, and capacity. Bars indicate mean and error bars show standard deviation. N=3; independent biological replicates are represented in blue, green, or pink colours. Black asterisks compare within a single treatment group; red asterisks compare across different treatments. **** P ≤ 0.0001, *** P ≤ 0.001, ** P ≤ 0.01.

In contrast to controls or inactive LAAO, both basal and maximal respiration levels were severely reduced by LAAO^WT^ treatment and correlated with high levels of proton leakage (Fig. 6C). During the time course investigated, there were fluctuations in ATP production and spare capacity in controls or catalytic inactive LAAO, but these were not significant. In contrast, both ATP generation and spare capacity were severely decreased by LAAO^WT^ incubation (Fig. 6D). However, non-mitochondria respiration (Fig. 6D) and glycolysis (Fig. 6E) were not altered by either treatment. Instead, a strong decline in glycolytic reserve levels, glycolytic capacity and non-glycolytic acidification were significantly reduced by LAAO^WT^ (Fig. 6E-F, Fig. S4).

Taken together, our data indicate that LAAO interferes with energy production and impairs the switch to alternative energy sources in a catalyses-dependent manner. The outcome is a significant reduction in respiration, glycolytic reserve and capacity and a shutdown of ATP production.

### LAAO induces mitophagy

The severe impairment of energy production and mitochondria morphology suggests that defective mitochondria could be cleared by mitophagy. However, we postulated that mitophagy may also be compromised, as the strong reduction in lysosome numbers and size may prevent autophagosome degradation (Fig. 3B-D). We evaluated the mitophagy index using live imaging and SH-SY5Y cells stable expressing mt-Keima (Fig. 7), a pH-sensitive fluorescent protein ^39^ that is targeted to the mitochondria. During mitophagy, in the acidic lysosomal environment, mt-Keima fluorescence emission shifts from the shorter (green) to longer wavelength (red). The mitophagy index was calculated as a ratio between the segmented area of the two mt-Keima in each image (lysosomal and cytoplasmic). As a positive control, hydrogen peroxide treatment to induce oxidative stress led to a progressive co-localization of green and red mt-Keima (Fig. S5A). In contrast, addition of oligomycin to inactivate mammalian F_0_F_1_-ATPsynthase did not promote mt-Keima shift to red fluorescence, consistent with the prevention of H^+^ ions transport and lysosome acidification (Fig. S5B).

**Fig. 7.**
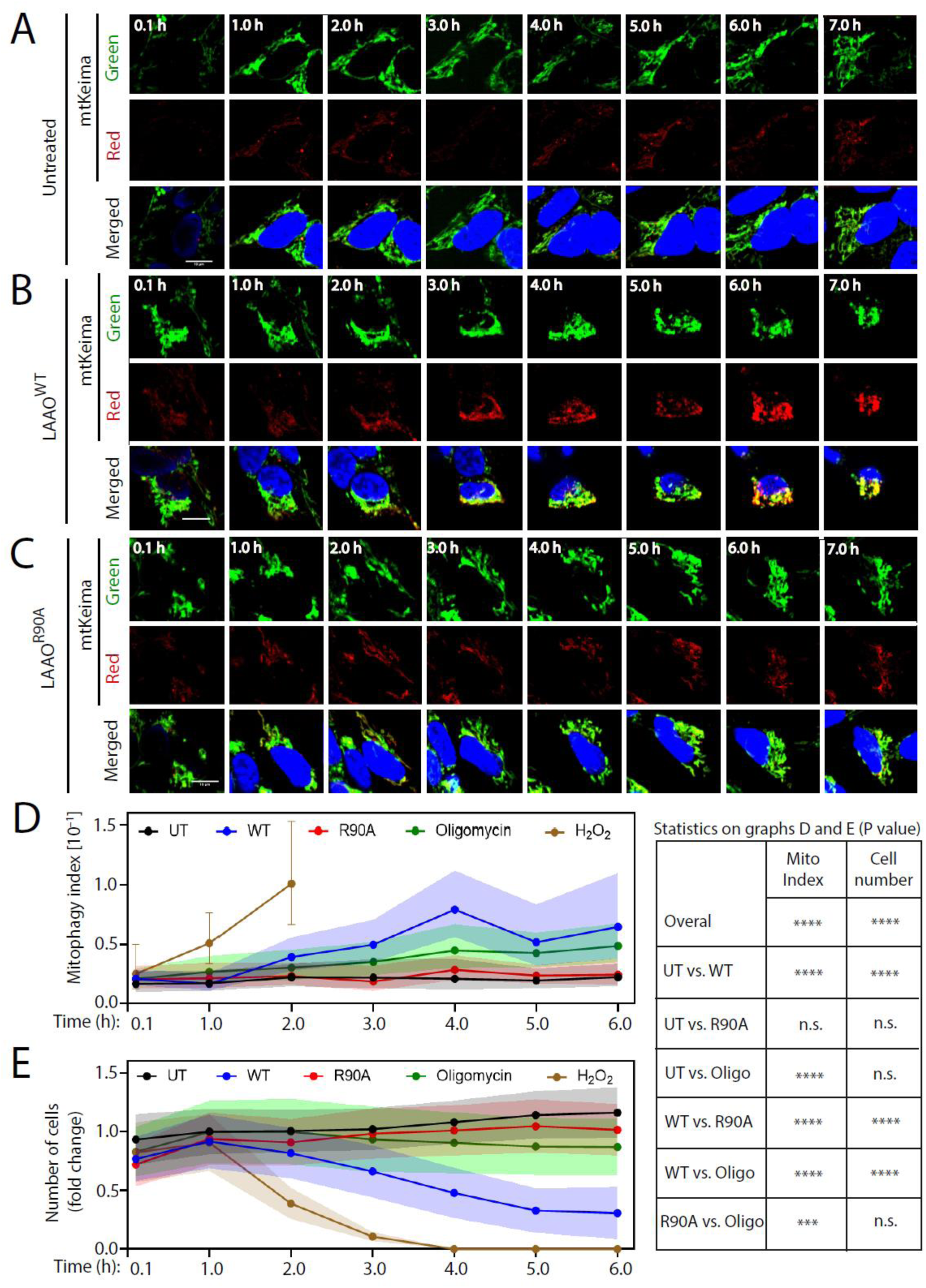
LAAO cytotoxicity triggers mitophagy, but not mitochondria clearance. **A-B,** Representative confocal live imaging of SH-SY5Y cells stable expressing mtKeima (mitochondria-targeted dual fluorescent protein) untreated (A) or following treatment with (B) LAAO^WT^ or (C) LAAO^R90A^ (2.8 X EC_50_ or 12.25 nM for SH-SY5Y cells). Images show fluorescence of mtKeima green (top row), mtKeima red (middle row), and the merged images (bottom row) including nuclei stained with Hoechst 33342 (at the lower concentration used, the nucleus cannot be fully stained at 0.1 h). As controls, cells were also treated with 10 µM oligomycin or 1 mM H_2_O_2_ (shown in Fig. S5). **D**, Mitophagy index assess the change of mitochondria state from cytoplasmic mtKeima (green at pH 7.0) to a lysosomal mtKeima (red at pH 4.0). Mitophagy index was calculated (mean area of red mtKeima/mean area of green mtKeima) per image and normalized to values of untreated (UT) samples. **E**, Number of cells present after treatment for up to 6 hours. Treatment with 1 mM H_2_O_2_ was toxic to SH-SYS5 cells and led to substantial cell detachment after 2-3 h. Values in D-E are shown as geometric mean with error, 95% confidence interval (95% CI) added as coloured shade or bar for H_2_O_2_. N=3. **** P ≤ 0.0001, *** P ≤ 0.001, ** P ≤ 0.01, * P ≤ 0.05. Table on the right shows P value for overall comparison. Scale bar: 10 µm.

The EC_50_ value of LAAO^WT^ for SH-SY5Y mt-Keima was determined (EC_50_ = 4.38 nM). Basal levels of mitophagy were similar in untreated samples and after incubation with inactive LAAO (LAAO^R90A^, Fig. 7A,C). LAAO^WT^ treatment promoted a persistent conversion to red emission from 2 hours onwards (Fig. 7B): the mitophagy index reached a maximum at 4 hours (Fig. 7D) and was significantly distinct from its inactive counterpart. There were no significant differences in mitophagy index over time between cells treated with LAAO^R90A^ or controls (Fig. 7D). Mitochondria degradation was not observed during the time course, due to the high sensitivity of SH-SY5Y cells to LAAO treatment.

No significant changes in cell numbers were observed in untreated, oligomycin-or inactive LAAO-treated samples (Fig. 7E). There was a sharp increase in mitophagy index and decrease in the number of cells treated with H_2_O_2_ after 2 hours (Fig. 7D-E) and at 4 hours. There was a 50% reduction in cell numbers in LAAO^WT^ samples (Fig. 7E). Thus, when treated with wild-type LAAO, a smaller number of cells have a very high mitophagy index. We surmise that mitophagy is a cellular response to counteract the impairment of mitochondria function by LAAO, and thus elicit elimination via organelle quality control.

All results put together (Fig. 8) showed a temporal segregation of the various cellular events. Despite normalization of LAAO EC_50_ for each cell type and condition when performing the experiments, distinct assay sensitivities and cell type-specific properties may affect the timing when a change in cellular events is detected. Nevertheless, a comparative analysis of phenotypes in a single cell type is feasible. Furthermore, while the timeframe might shift, key results confirmed among cell lines indicate the phenotype reproducibility across different cell types.

**Fig. 8.**
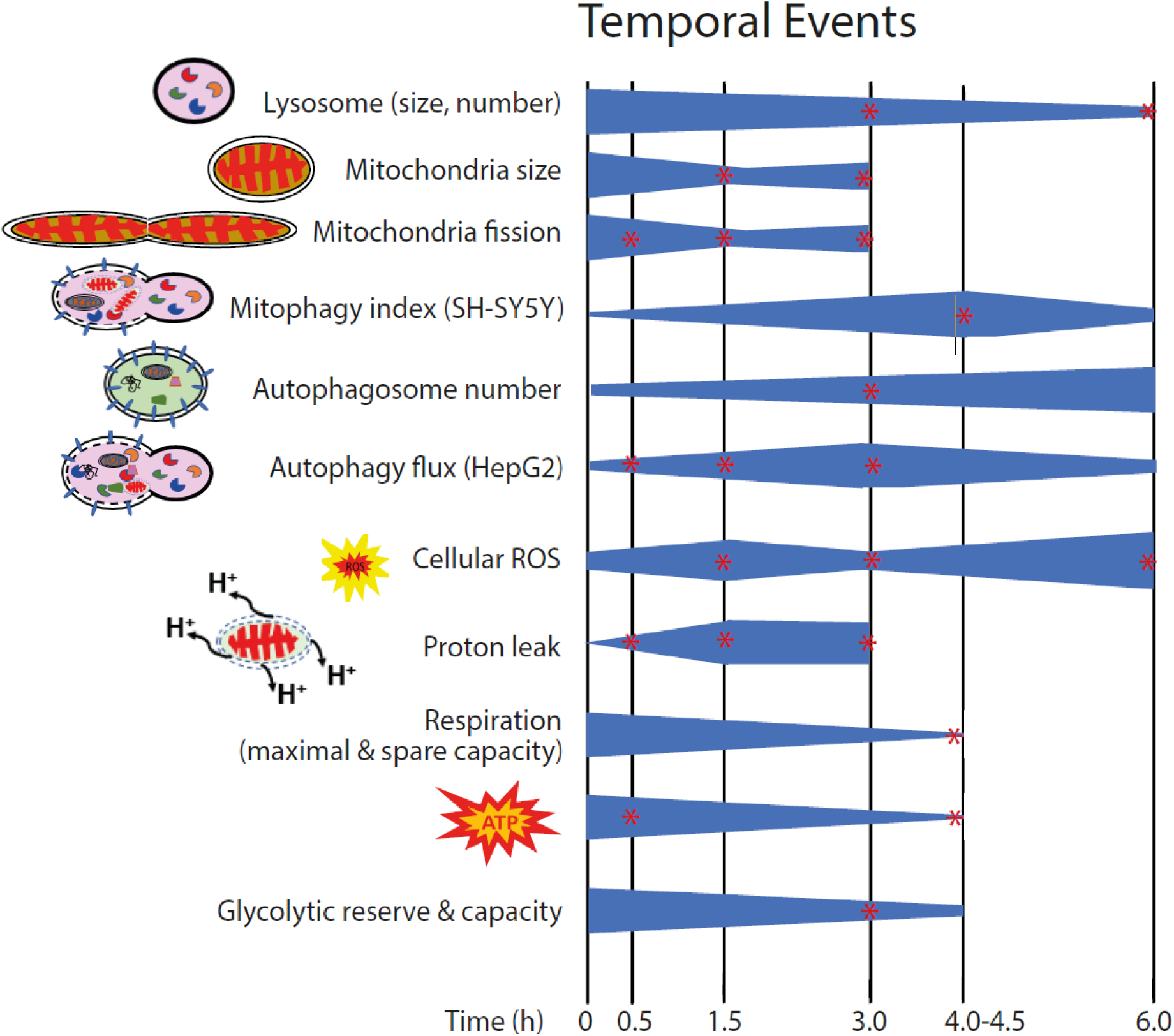
Summary of temporal events triggered by treatment with *B. atrox* LAAO^WT^. Mapping the changes of various cellular processes during LAAO envenomation. Quantitative morphometry was used to assess: (i) size and numbers of mitochondria and lysosomes (Fig. 3-5) and (ii) autophagosome numbers (Fig. 2). Functional assays evaluated autophagy flux index (Fig. 2), mitophagy index (Fig. 7), oxidative stress species (cellular ROS (Fig. 1). Mitochondrial functions and energy generation (Fig.6) are shown as proton leak, mitochondria respiration (maximal and spare capacity), ATP production, and glycolytic capacity and reserve. Normal keratinocytes were used in most assays. HepG2 cells expressing tandem mCherry-EGFP-LC3 and SH-SY5Y cells expressing mt-Keima were used as well-established models for autophagy flux index and mitophagy index, respectively. The symbol * denotes statistical significance and p values can be found in the respective figures.

## DISCUSSION

Here we contribute to the conceptual understanding of the cellular events leading to tissue dysfunction triggered by a snake venom toxin, LAAO. Here, we identify unappreciated cellular processes disrupted by LAAO prior to induction of cell death and necrosis ^15^. The reduced cell fitness induced by LAAO is catalysis-dependent and results from a coordinated damage in mitochondria and lysosomes (perturbed organelle morphology, function, and clearance), resulting in the shut-down of cellular metabolism and energy production.

The use of recombinant LAAO (glycosylated, soluble, and active) demonstrates unequivocally the relative contribution of essential residues for its enzymatic catalysis, which has been controversial in the past ^29,31,40,41^. Mutation R90A or Y372A strongly impairs LAAO^WT^ activity *in vitro*, consistent with the prediction of R90 as a co-factor (FAD) binding and cooperation with Y372 in substrate interaction. Glycosylation at residue N172 has been indirectly shown as relevant for catalysis ^27^. LAAO^N172A^ mutant is indeed inactive, which may result from N-linked glycosylation contribution to maintain protein stability ^42^ or to regulate the exit of H_2_O_2_ from the catalytic pocket ^29^. Thus, LAAO residues R90, N172 and Y372 have key catalytic functions, whereas other proposed amino acids are not essential (H223) or reduce (R322) activity. The latter two residues do not have a well-established role ^29,31^.

Recombinant LAAO^WT^ cellular phenotypes reproduce those observed with native LAAO treatment ^15^: higher levels of reactive oxygen species (ROS) and number of autophagosomes. We think it is unlikely that the higher autophagosome levels result from oxidative stress or amino-acid modification or catabolism ^24^, as both wild-type and catalytic dead LAAO increase autophagosome numbers. Instead, we demonstrate that LAAO^WT^ attenuates autophagy flux index (i.e., autophagosome clearance) in a catalysis-dependent manner. An adaptative cellular response to oxidative stress generated by LAAO may promote autophagy flux via (i) altered catabolism of amino acids necessary to produce antioxidants (e.g., glutamine) or (ii) unbalancing the homeostasis of LAAO amino acid substrates, some of which (i.e., leucine) are key regulators of the amino acid sensing machinery ^24^. Furthermore, the by-products of LAAO-dependent oxidation (H_2_O_2_ and ammonia) can themselves contribute to lysosomal defects ^23,43^ and potentially impair cargo degradation, lysosome acidification and/or fusion with autophagosomes.

First, LAAO^WT^ treatment strongly decreases lysosome size and number, which limits availability of degradative enzymes and reduces membrane area for appropriate regulation of amino acid sensing by mTORC1 complexes ^44^. Second, internalised LAAO^WT^ partially co-localizes with lysosomes, indicating that LAAO is at the right place to participate in lysosomal dysfunction. Such co-localization is novel for a snake toxin, and consistent with the preferential localization of its homologue IL4I1, a mouse B lymphocyte LAAO ^45^. At the lysosomal acidic compartment, both enzymes have the optimal pH for their activity ^45^. As lysosomes also participate in signalling, exocytosis, and nutrient sensing ^46,47^, extensive lysosomal impairment by LAAO may also substantially damage cell homeostasis.

In addition to degrative processes, we find that LAAO^WT^ cytotoxicity (i) impairs mitochondria dynamics, the coordinated fission, and fusion events ^48,49^ and (ii) alters mitochondrial shape, size, and distribution in keratinocytes. Interference with mitochondrial function appears to be a main feature of native LAAOs from different snake venoms (altered mitochondrial membrane potential and Cytochrome C release) ^14,35,50^. Furthermore, modulation of mitochondrial dynamics by oxidative stress and amino acids depletion is observed in other model systems ^51^, and likely to contribute to LAAO-induced mitochondrial defects. Thus, LAAO interferes with fundamental cellular processes that have a domino effect in shutting down cell viability.

We predict that the impairment of mitochondrial dynamics by LAAO may also damage energy generation and consumption ^52^. The mitochondrial network is highly dynamic and intimately associated with optimal bioenergetic capacity and the regulation of cellular signalling and metabolism ^20^. The strong perturbation of mitochondria dynamics by LAAO^WT^ restricts cell recovery. The rate of glycolysis and non-mitochondrial respiration over time are similar in controls and LAAO^R90A^. In contrast, LAAO^WT^ strongly reduces basal respiration and maximal mitochondrial capacity, thereby limiting ATP production. Under these conditions, cells reduce their ability to respond to energetic demand (spare capacity depletion) and adapt metabolically by using their glycolytic reserve to maintain glycolysis rate unchanged. In addition, LAAO^WT^ treatment prevents switching energy sources (glycolysis to oxidative phosphorylation) and reduces non-glycolytic acidification. Therefore, LAAO^WT^ impairs an adaptive energetic response that would facilitate cell recovery from mitochondria dysfunction.

Oxidative stress may trigger mitochondrial fission and mitophagy ^20,53^, and indeed mitophagy is stimulated by LAAO^WT^ exposure to remove damaged and smaller sized mitochondria. In mt-Keima-expressing SH-SY5Y cells, fusion of mitochondria-containing autophagosomes with lysosomes occurs, but not their degradation. The attenuated clearance of mitochondria in SH-SY5Y cells in the presence of LAAO^WT^ is consistent with the inhibition of autophagy flux shown in HepG2 cells. The reproducibility of the lysosomal and mitochondrial defects in different cell types is significant for our understanding of the envenomation process.

In summary, our work reveals unprecedented molecular and cellular mechanisms triggered by snake venom LAAO (Fig. 8). The mapped cellular phenotypes are temporally distinct, cooperatively reduce cell survival, and are likely to be interdependent. In the keratinocyte model, higher levels of proton leakage, and increased mitochondria fission are the first detected events after LAAO treatment in keratinocytes, followed by increased oxidative stress. Later, an increase in the number of autophagosomes coincides with the reduced lysosome size and numbers. Such lysosomal defects contribute to impairment of organelle quality control and clearance, via inhibition of autophagy flux and mitophagy in different cell models (HepG2 and SH-SY5Y models, respectively). Alongside these phenotypes, there are profound changes in metabolic rates, energy production and consumption, which further compromise cell fitness. Thus, the inability to degrade and recycle damaged organelles in the presence of LAAO^WT^ is highly likely to synergize with oxidative stress and energetic impairment during cytotoxicity.

Overall, our study provides a framework that is instrumental to enhance the knowledge of the envenomation process. The early, multiple organelle impairment caused by LAAO exposure highlights points of future therapeutic intervention to restore tissue homeostasis and renewal in an envenomation context.

## ACKNOWLEDGMENTS

We thank Fernanda Costal-Oliveira for technical support. K.P. was supported by the Wellcome Trust and Global Challenges Research Fund (GCRF)-England. J.K.V. and R.K. were supported by Medical Research Council core funding from the MRC-UCL University Unit Grant (Ref MC_U12266B) and the MRC Dementia Platform UK (MR/M02492X/1). CC-O was supported by Conselho Nacional de Desenvolvimento Científico e Tecnológico (CNPq), Coordenação de Aperfeiçoamento de Pessoal de Nível Superior (CAPES) and Fundação de Amparo à Pesquisa do Estado de Minas Gerais (FAPEMIG)-Brazil.

## AUTHOR CONTRIBUTIONS

K.P. performed most of the experiments; C.L. performed mitophagy experiments and interpreted the data on that section. E.T. contributed to the cell Mito stress test experiments. J.K.V. conducted confocal imaging and image analysis. V.B and K.P. conceived the study and wrote the manuscript. V.B. supervised the research and secured funding. R.K. helped with the conception and design of the project and together with C-CO edited the manuscript. All authors reviewed the manuscript.

## METHODS

### Plasmid cloning

Wild-type LAAO from *B. atrox* (LAAO^WT^) was inserted into pcDNA3.1 vector with a His-tag at the C-terminus as described elsewhere (manuscript in preparation). To create point mutations in pcDNA-LAAO^WT^, relevant primers were designed (Table S1) to introduce the following mutations: R90A (LAAO^R90A^), N172A (LAAO^N172A^), H223A (LAAO^H223A^), R322A (LAAO^R322A^) and Y372A (LAAO^Y372A^). We aimed to test the following predictions: R90A mutation is predicted to remove the interaction between LAAO, substrate and co-factor FAD, while N172A abolishes a glycosylation site. H223A and R322A mutations are predicted to interfere with substrate binding sites and Y372A should interfere with both substrate/ ligand binding site. All PCRs were conducted for 30 cycles using Pyrobest DNA polymerase (Takara, R005A), PCR products were phosphorylated and ligated. All clones were verified by DNA sequencing.

### Cell culture

All cells were cultured in a humid chamber at 37°C and 5% CO_2_. HEK293T (human embryonic kidney cells) were used to produce recombinant LAAO proteins only. They were grown in a 500 cm^2^ large Nunc™ square culture dish (Thermo Scientific, 240835; 100 ml culture media) in DMEM medium containing high glucose (Sigma, D6546) supplemented with GlutaMax (Gibco, 35050-061) and 10% foetal calf serum (FCS; Seralab, EU-000-F) until they reach the confluency of around 80%. The cells were then transfected with 50 µg of plasmids pcDNA-LAAO, pcDNA-LAAO^WT^, pcDNA-LAAO^R90A^, pcDNA-LAAO^N172A^, pcDNA-LAAO^H223A^, pcDNA-LAAO^R322A^ or pcDNA-LAAO^Y372A^ using FuGENE® HD transfection reagent at 1 µg plasmid / 4 µl transfection reagent (Promega, E2312).

Normal human epidermal keratinocytes at 4000 cells/well (passages 3-6) were co-cultured with 3T3-J2 fibroblasts in a 96 well plate (Corning, 3585; cell viability assay) or a CellCarrier-96 ultra microplates (PerkinElmer, 6055300; confocal imaging) in FAD standard medium (FAD medium (Gibco, custom made) as described previously ^54^). Where indicated, cells were transfected with pEGFP-LC3 (EGFP-LC3: microtubule associated protein 1 light chain 3 fused with enhanced green fluorescent protein) (Addgene, 24920) at 0.1 µg of the plasmid (in 50 µl Opti-MEM® Reduced Serum Media) and 0.4 µl of FuGENE® HD Transfection Reagent.

To investigate autophagy flux and mitophagy, two different cell models were used that are well established in the field: HepG2 (human liver cancer cell line) stably expressing mCherry-EGFP-LC3 and SH-SY5Y (neuroblastoma cell line) cells stably expressing mitochondria-targeted monomeric Keima (mtKeima, (MBL, AM-V0251HM) fluorescent protein, respectively. Cells were seeded at 10000 cells/well in a 384 well plate (Nunc, 164688; cell viability assay or a CellCarrier-384 ultra microplates (PerkinElmer, 6057300; confocal imaging) in Dulbecco’s modified Eagle’s medium containing GlutaMax (Gibco, 61965-026) supplemented with 10% FCS.

### Recombinant protein production and purification

After transfection for 4 hours, the medium was refreshed with medium containing 10 µg / ml catalase (Sigma, C1345) to initiate collection of recombinant protein produced by the HEK293T secretion system. Medium was collected every 24 h and replaced with fresh medium as described above. After 3 days, the collected media was concentrated 10-fold using Amicon Ultra centrifugal filter units, Ultra-15, MWCO 10 kDa (Sigma, Z706345-8EA). Imidazole at 20 mM (Sigma, 792527-100G) and 1 mM dithiothreitol were added, and the secreted medium was incubated with 1.5 ml of Ni-NTA Agarose (Qiagen, 30210) for 1 h shaking at 4 °C for the isolation of recombinant His-tagged LAAO. Beads were centrifuged and washed 5 times with PBS buffer pH 7.3 (140 mM NaCl, 2.7 mM KCl, 10 mM Na2HPO4, 1.8 mM KH2PO4) containing 20 mM imidazole and 1 mM DTT. LAAO^WT^ and mutants were eluted three times with 2 ml PBS buffer pH 7.3 containing 150 mM imidazole and 1 mM DTT. The proteins were concentrated to 1 ml and washed with 50 mM sodium acetate buffer pH 5.0 containing 150mM NaCl and 1 mM DTT. For long-term storage, glycerol was added (10% volume/volume) and the proteins were aliquoted and stored at −80°C.

Protein concentrations were determined by loading different amounts of recombinant LAAO on a 10% sodium dodecyl sulphate (SDS)−polyacrylamide gel electrophoresis and staining with Coomassie Blue. Different concentrations of bovine serum albumin (BSA) (Thermo Fisher, 23209) were loaded as a standard in each gel. Bands intensities were quantified using Fiji and the apparent concentration of recombinant LAAO derived from the corresponding BSA standard curve.

### Activity assays and enzyme kinetics of LAAO^WT^ and mutants

Enzymatic assay for LAAO activity was conducted with modifications from what described ^55^. To determine LAAO activity in the secreted media, 30 µl of secreted medium and 70 µl of LAAO assay solution (0.1 M Tris-HCl pH 8.0, 5 mM L-leucine, 2 mM OPD (o-phenylenediamine dihydrochloride) (Sigma, P9187), 33.3 µg/ml peroxidase (Sigma, 77332)), were added to 96 well ELISA plate (Thermo Fisher, 612U96) and incubated for 1 h at 37°C. The reaction was stopped by adding 50 µl of 1M of sulphuric acid (H_2_SO_4_). The absorbance was determined at 490 nm using a SpectraMax i3X and Pro 6.5 software.

To investigate the activity of purified recombinant LAAO^WT^ and mutants in the presence of excessive substrate L-leucine (5 mM), 52.72 nM (or 0.3 µg per 100 µl) of the enzyme was used. The assay was done as describe above except that the purified protein (0.3 µg protein in 10 µl buffer (0.1 M NaCl, 50 mM CH_3_COONa pH 5.0)) was mixed with 90 µl of LAAO assay solution instead of 70 µl.

To determine enzyme kinetics, purified LAAO^WT^, LAAO^H223A^ and LAAO^R322A^ at 26.36 nM and LAAO^R90A^, LAAO^N172A^ and LAAO^Y372A^ at 158 nM were incubated with 2-fold serial dilutions of L-leucine from 10 to 1280 µM in an LAAO assay solution (0. 1M Tris-HCl pH 8.0, 2 mM OPD, 33.3 µg / ml peroxidase) in a 96 well ELISA plate (100 µl final assay volume). Samples were incubated at 37 °C for 1 h for LAAO^WT^, LAAO^H223A^ and LAAO^R322A^ or for 2 h for LAAO^R90A^, LAAO^N172A^ and LAAO^Y372A^ (mutants predicted to be inactive). The reaction was stopped by adding 50 µl of 1 M H_2_SO_4_ and the absorbance was determined at 490 nm as stated above. The initial velocity (µM per second) was defined as the change in the concentration of H_2_O_2_ produced by LAAO, which was plotted versus the concentration of substrate L-leucine before reaction (µM). The curves were then fitted using the nonlinear regression method in R software, from which the V_max_ and K_m_ (Michaelis constant) for each enzyme’substrate reaction was derived. The k_cat_ (catalytic constant) was determined by dividing V_max_ by the enzyme concentration. The catalytic efficiency is defined as K_cat_/ K_m_ (inverse µM per second).

### Determination of H_2_O_2_ production

To monitor the amount of H_2_O_2_ produced, 28 nM of LAAO^WT^ was added to DMEM phenol-free (Sigma, D1145) supplemented with 1% FCS and Glutamax and incubated at 37 °C, 5% CO_2_ for a period from zero to 6 h. Samples (50 µl) were collected to a 96-well ELISA plate (Thermo Fisher, 612U96) and mixed with 50 µl solution (4 mM OPD and 66.7 µg/ml peroxidase) for 5 min. The reaction was stopped by adding 50 µl of 0.5 M H_2_SO_4_. The absorbance was determined at 490 nm using a SpectraMax i3X and Pro 6.5 software. The amount of H_2_O_2_ content in the samples were calculated using a H_2_O_2_ standard curve.

### Cytotoxicity assay

Different cells show distinct sensitivity to LAAO toxicity ^35^; thus, the Effective Concentration to reduce viability to 50% in 24 hours (EC_50_) was calculated for each cell type (keratinocytes, HepG2 and SH-SY5Y) and batches of recombinant LAAO. LAAO cytotoxicity was tested using Alamar Blue cell viability reagent (Thermo Fisher, DAL1100) and the assay was performed as previous described ^56^ with modifications. After reaching 50-60% confluence, keratinocytes were treated with 2-fold serial dilutions from 7.5 to 0.00078125 µg/ml of freshly purified LAAO^WT^ in FAD standard medium containing 1% FCS for 24 h. The medium was then replaced with DMEM phenol-free (Gibco, 31053-028) containing 1% FCS, 4 mM L-glutamine and 10% Alamar Blue solution. Cells were continued to culture for 3 h before the fluorescence was read at 560 nm of excitation and 590 nm of emission in a POLARstar Galaxy fluorimeter using FLUOstar Galaxy software. For the comparison of cytotoxicity of LAAO^WT^ and mutants at the EC_50_ value of the WT, 14 nM of the WT and mutants were used to treat keratinocytes. For metabolic assays, normal keratinocytes were grown at higher density and thus a much higher EC_50_ was used in these experiments.

For the cytotoxicity assay of LAAO in SH-SY5Y or HepG2, cells were grown for 24 h before the treatment with LAAO^WT^ at 2-fold serial dilutions from 1.2 to 0.05 µg/ml and 7.0 to 0.5 µg/ml, respectively, for 24 h and processed as described above. Cytotoxicity at different conditions was calculated comparing to control set as 100% survival.

### Autophagosomes, organelles and oxidative stress measurements

After transfection with pEGFP-LC3 for 4 hours, keratinocyte medium was refreshed with a fresh FAD standard medium containing 10% FCS. After 48 h post-transfection, the culture medium was changed to FAD standard medium containing 1% FCS and treated with LAAO^WT^ or mutant LAAO^R90A^ at 28 nM for 0.5, 1.5, 3 and 6 h and stained live with 100 nM Lysotracker red DND-99 (Thermo Fisher, L7528) for 1.5 h before ending the treatment. The probe is a red-fluorescent dye that is highly selective for labelling and tracking acidic organelles.

For mitochondrial membrane potential assay at 2X EC_50_, cells were treated with 28 nM LAAO^WT^ for 0.5, 1.5, 3, and 6 h. To analyse the effect of LAAO^WT^ and catalytic dead mutant R90A on mitochondrial membrane potential, mitochondrial size and Feret’s diameter, ROS generation, 280 nM of LAAO^WT^ or LAAO^R90A^ was used for different time points. Cells were stained live with 100 nM MitoTracker Red CMXRos (Thermo Fisher, M7512), 5 µM CellRox Green Reagent (Thermo Fisher, C10444) and 1:10000 Hoechst 33342 for 1.5 h. CellRox Green Reagent is a fluorogenic probe; the dye is weakly fluorescent while in a reduced state and exhibits bright green photostable fluorescence upon oxidation by reactive oxygen species and subsequent binding to DNA.

After live labelling with LysoTracker, MitoTracker Red CMXRos, or CellRox Green as described above, cells were washed once with PBS and fixed with 4% PFA (Alfa Aesar, J61899) for 10 min and stored in PBS at 4°C when necessary. Images were captured at 40x for 20 field of views/well of a 96 well plate.

### Monitoring autophagy flux

HepG2 cells stably expressing tandem mCherry-EGFP-LC3 were used to monitor autophagy flux ^33^ via a live-cell imaging based approach. Cells were seeded in CellCarrier-384 ultra microplates (PerkinElmer, 6057300) at 10000 cell/well/30 µl. After 16 h, the media of the culture was replaced with DMEM + GlutaMax supplemented with 10% FCS and 1:16000 Hoechst 33342. Cells were left untreated or treated with 42 nM (2.4 µg/ml) LAAO^WT^ or LAAO^R90A^, 10 µM chloroquine in the presence or absence of 42 nM LAAO^WT^. Images were captured at 40x in several fields of view and the number of LC3 puncta under mCherry or EGFP channel were calculated, and the autophagy flux index was determined. The autophagy flux index (in each field of view) = total number of LC3 puncta in mCherry channel/total number of LC3 puncta in EGFP channel.

### Bioenergetic measurements

In order to measure glycolysis and mitochondrial respiration, in real time, the Agilent Seahorse XFe96 Extracellular Flux Analyzer (Seahorse Biosciences) was used, as described previously ^57^. The outputs were recorded as ECAR and OCR, respectively. In brief, keratinocytes were co-cultured with J2 in 9 cm dish until they reach around 80% confluency. J2 cells were removed from the culture dish by briefly washing with Versene. Keratinocytes were seeded to a 1 μg/well collagen I (Sigma, C3867-1VL) precoated 96 well plate (Agilent, 102601-100) at a density of 19000 cells/well/80 μl. The cells were rested at RT for 45 min before transferring to incubator. After 16 h, the culture medium was changed to FAD standard medium containing 1% FCS and treated with 73 nM LAAO^WT^ or LAAO^R90A^ for 0.5, 1.5. 3.0, and 4.5 h. Culture media was then replaced with 175 μL of XF DMEM (Agilent, 103575-100) media containing 10 mM glucose, 1 mM sodium pyruvate, and 2 mM L-glutamine for Mito stress test or with 180 μL of the XF DMEM media containing only 2 mM L-glutamine for Glycolysis stress test. Cells were then incubated for 45 min at 37 °C without carbon dioxide to allow for cells to reach the ideal pH and temperature conditions required for the assay. The machine ran a 3-minute mix and 3-minute read cycle which generated OCR and ECAR readings. During the assay, various compounds were injected via ports to see their effects on mitochondrial respiration and glycolysis. Three OCR measurements were recorded after each port injection starting with oligomycin (1.5 μM), followed by FCCP (1 μM), and lastly a combination of rotenone and antimycin A (1 μM each) including Hoechst 33342 (1:10000). ECAR measurements were recorded similarly except the port injection started with glucose (10 mM), followed by oligomycin (2 μM), and lastly 2DG (50 mM) including Hoechst 33342 (1:10000). The final volume of each assay well became 250 μL. After analysis by the XFe96 Extracellular Flux Analyzer, the plate was imaged in the Opera Phenix (PerkinElmer) for capturing the image of the nucleus. Values of the 3^rd^ reading was used for the calculation of mitochondrial activity and glycolytic activity parameters and calculated based on the Agilent Seahorse guidance.

### Detection of mitophagy using SH-SY5Y cells stable expressing mtKeima

SH-SY5Y stably expressing mtKeima cells were used to assess mitophagy activity induced by LAAO, hydrogen peroxide, and oligomycin in live cells as the fluorescent of the Keima is visualized green (cytoplasmic mitochondria) or red (lysosomal mitochondria) at pH7 or pH4, respectively ^39^. After 16 h of seeding cells, the media was replaced with DMEM supplemented with GlutaMax, 10% FCS and 1:16000 Hoechst 33342. Cells were treated with 12.25 nM (0.7 µg/ml) LAAO^WT^, 12.25 nM LAAO^R90A^, 1 mM H_2_O_2_, or 10 µM oligomycin. Live images were captured at 63x for 9 field of views/well of a 384 well plate at 0.1 h and every hour from 1 to 6 h. The images were analysed using Columbus 2.8 (PerkinElmer) and the mitophagy index was determined. Briefly, cells and mtKeima were segmented and the area of cytoplasmic mtKeima (green mtKeima; cytoplasmic mitochondria) and lysosomal mtKeima (red mtKeima; mitochondria in lysosomes) was determined. The mitophagy index (in each field of view) = total area of lysosomal mitochondria / total area of cytoplasmic mitochondria.

### Immunocytochemistry and confocal microscopy image acquisition

After fixing with 4% PFA, keratinocytes were washed 3 times and permeabilized with 0.1% Triton X-100 and 10% FCS in PBS for 10 min and block with 50% FCS in PBS for 1 h at RT after 3 washings. Cells were incubated with anti-6x His-tagged antibody (DyLight 650) at 1:2000 (Abcam, ab117504) for 1 h or with anti-6x His-tagged antibody (Abcam, ab18184) at 1:1000 for 1 h followed by Cy5 secondary antibody (Jackson Immuno Research) at 1:1000 for 1 h for the localization of LAAO. Cells were finally stained with 1:10000 Hoechst 33342 (Thermo Fisher, H3570) for 10 min and were covered with 100 µl PBS. They were imaged at 20 images/well of a 96 well plate at 40× (pixel size 0.1494 µm) using Opera Phenix High Content Screening System (PerkinElmer) with water immersion NA = 1.1.

Live images cells were recorded in a humid chamber at 37 °C and 5% CO_2_ at 40× (pixel size 0.1494 µm) or 63× (pixel size 0.0949 µm) using Opera Phenix High Content Screening System (PerkinElmer). Excitation and emission wavelengths were adjusted based on manufacturer instructions.

### Image analysis

Image processing was done on a SuperServer 4048B-TR4FT high-performance server grade barebone computer equipped with 4 eight-core Intel Xeon E7-4809 v3 CPUs and 1 TB RAM running Ubuntu Linux 14.04 LTS. For desktop computation, a custom workstation was used with specifications Intel Core i9-7900X CPU and 128GB RAM memory running 64-bit Windows 10 Pro operating system.

Image processing was performed using Fiji/ImageJ ^58^ version 1.52n and Java 1.8.0_172. Images were acquired with Opera Phenix (Perkin Elmer) and exported with Harmony 4.8 software (Perkin Elmer) as single channel 16-bit TIF images.

The number of nuclei were calculated by pre-processing the nuclear channel images with a 20-pixel radius mean filter followed by a watershed segmentation with a specified noise tolerance/prominence parameter. The nuclei area and ROS intensity was quantified by segmenting the nuclear channel images using a manually selected threshold and measuring both the area of the binary nuclear mask and the mean intensity of the underlying ROS channel pixels within the nuclear mask. The number of LC3 puncta was quantified by segmenting the LC3 images by watershed segmentation with suitably chosen noise tolerance/prominence parameter.

Lysosome images were pre-processed by contrast stretching subsequentially converted to 8-bit depth. Auto local segmentation was performed by Bernsen’s thresholding method (https://imagej.net/Auto_Local_Threshold) using parameters 5-pixel radius for local domain size and 30 for contrast threshold. Lysosome number and size (area) were measured on the segmented binary image using the Particle Analysis function after applying a size filter that kept only the particles larger than 0.05 µm^2^.

Mitochondria membrane potential was calculated as mean intensity of the membrane potential channel pixels under the binary mask of tissue which is calculated as follows. Images were pre-processed by contrast stretching followed by conversion to 8-bit depth. A mean filter with 50 pixels radius was applied and finally a binary mask was generated by thresholding with the intensity value of 5.

Mitochondria were locally segmented by Bernsen’s thresholding method in 5-pixel radius for local domain size and default value for contrast threshold. The mitochondrial number, area and Feret’s diameter, which represents the longest distance (μm) between any two points within a given mitochondrion. Values were quantified for each mitochondrial fragment with the size filter that only the particles larger than 0.05 µm^2^ were kept.

### Statistical analysis

All statistical tests were performed using GraphPad Prism. For comparison of data from many groups, one-way analysis of variance was performed using Kruskal-Wallis and Dunn’s multiple comparison test to determine statistical significance. For experiments with time dependent recorded from live imaging with multiple treatments, two-way analysis of variance was performed using Tukey’s multiple comparison test to determine statistical significance. The number of images analysed in each treatment per replicate/experiment is shown in File S1.

## AVAILABILITY OF DATA AND MATERIALS

All data supporting the findings of this study are available within the paper and its supplementary information.

---

**Fig. S1.**
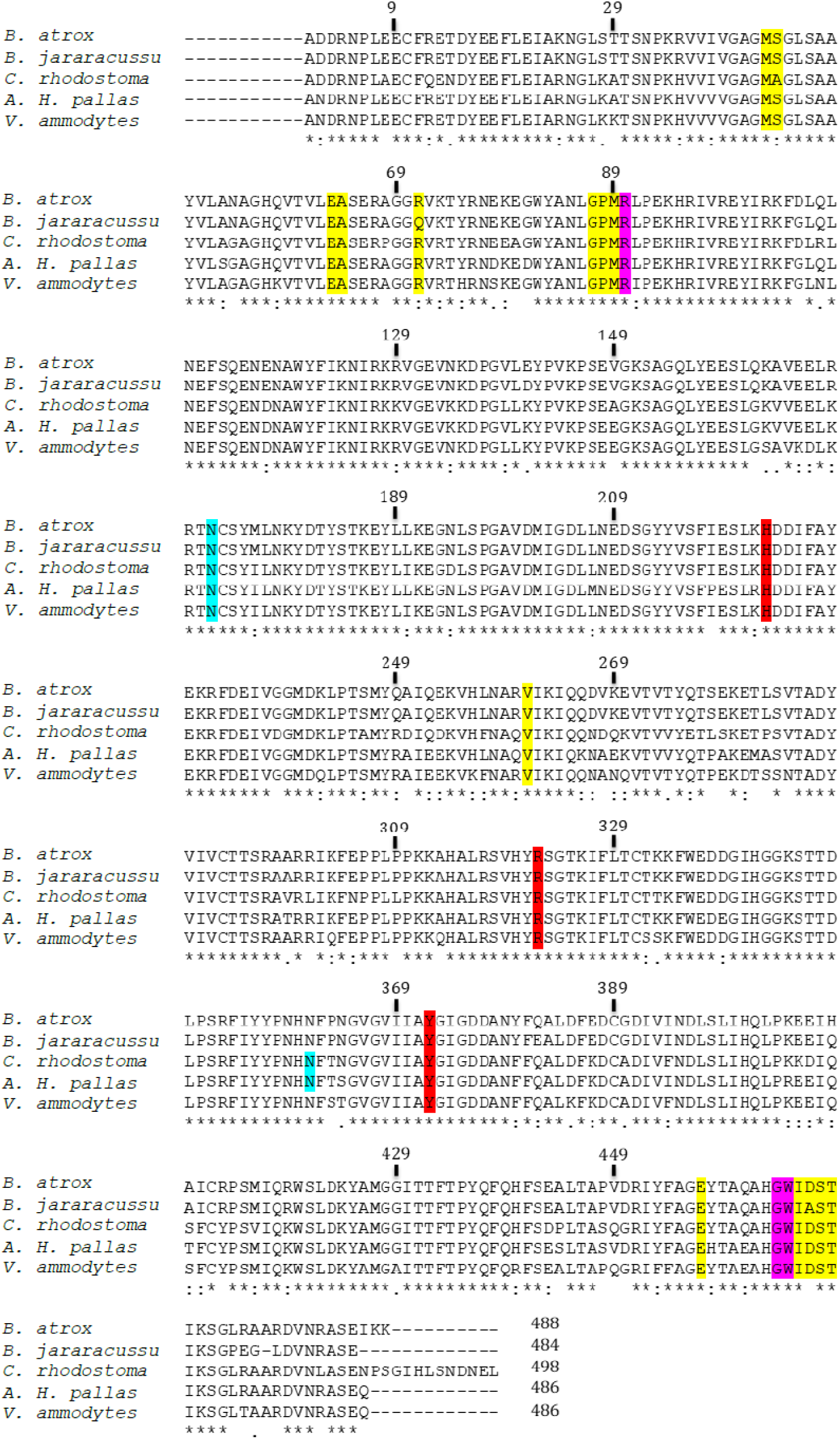
Alignment of the amino acid sequences of known snake venom LAAO structures. LAAO sequences from the species *B. atrox* (5TS5)*, B. jararacussu* (PDB 4E0V), *C. rhodostoma* (PDB 2IID)*, A. H. pallas* (PDB 1REO)*, and V. ammodytes* (PDB 3KVE) were aligned using Clustal Omega (1.2.1) multiple sequence alignment. Predicted cofactor FAD binding residues are shaded in yellow; and substrate binding residues are shaded in pink. Amino acid substitutions are denoted as: * (identical residues);: (conserved substitutions);. (semi-conserved substitutions).

**Fig. S2.**
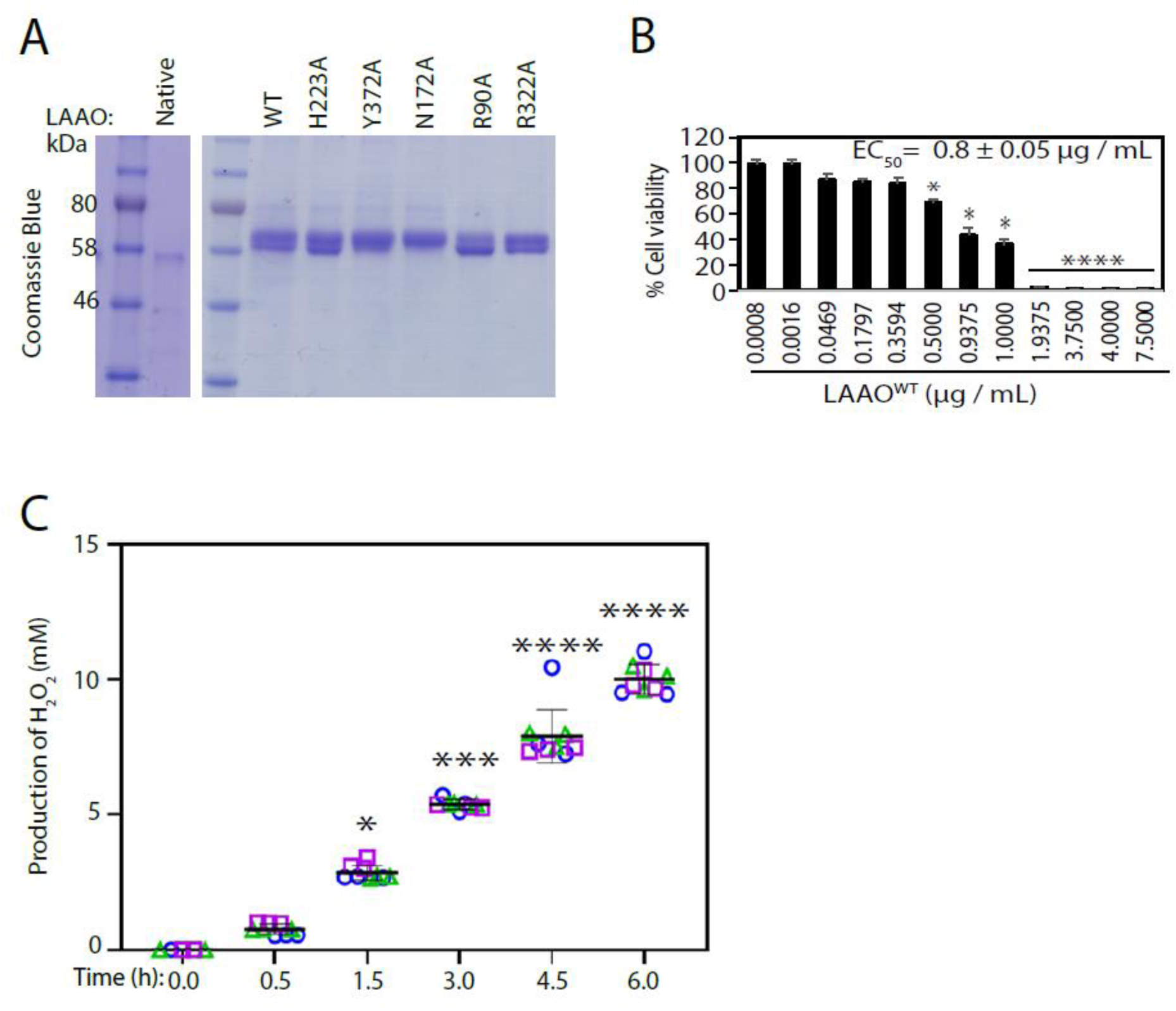
*B. atrox* LAAO^WT^ induces cell cytotoxicity. **A**, Recombinant LAAO-6xHis wild-type (WT) and mutants were purified, separated in SDS−PAGE and stained with Coomassie brilliant blue. Native LAAO purified from *B. atrox* venom was used as control. Purified recombinant LAAOs has a lower mobility than native LAAO (UniProtKB: P0CC17; 488 aa, approximately 55.5 kDa), due to additional amino acid residues for targeting for secretion and His-tag sequence. Glycosylation differences between mammalian (HEK293 cells) and reptilian cells may also contribute to its distinct electrophoretic mobility. **B**, Calculation of Effective concentration of LAAO^WT^ (EC_50_) in keratinocytes. Cells were incubated with different concentrations of LAAO for 24 h and the metabolic cell fitness was analysed using Alamar Blue. **C,** Determining the amount of H_2_O_2_ produced by LAAO^WT^ when added to the medium in the absence of cells. LAAO^WT^ (28 nM) was added to DMEM phenol-free supplemented with 1% FCS and Glutamax and incubated at 37 °C, 5% CO_2_ for a period from zero to 6 h. The amount of H_2_O_2_ content in the samples were calculated using a H_2_O_2_ standard curve. Results are displayed as mean ± standard deviation. N=3.

**Fig. S3.**
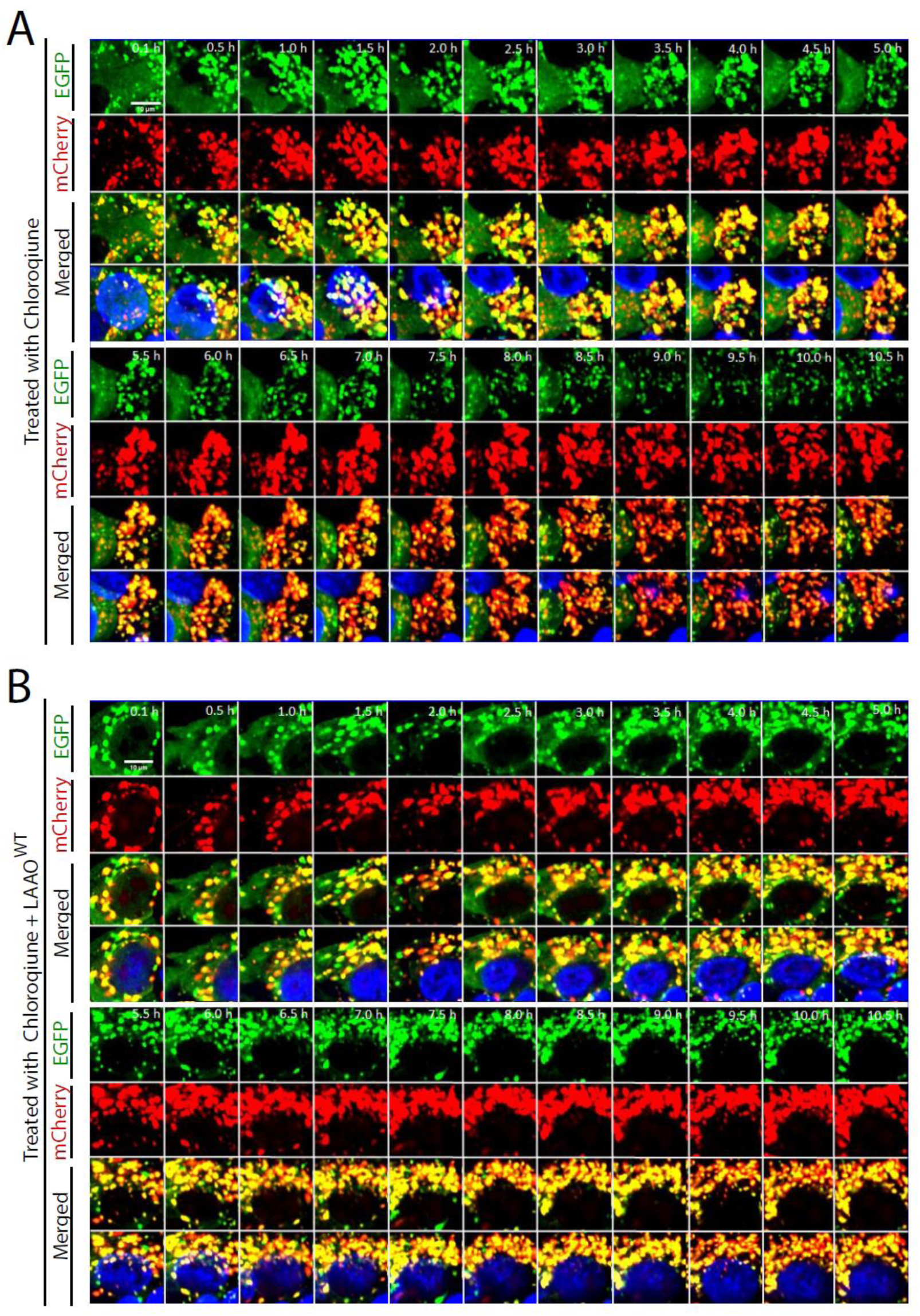
Responses of HepG2 cells stable expressing tandem mCherry-EGFP-LC3 to treatment with LAAO^WT^ during a time course in the presence or absence of chloroquine. HepG2 cells stable expressing tandem mCherry-EGFP-LC3 were treated with 10 µM of chloroquine or 10 µM of chloroquine and 1X EC50 (42 nM for HeG2 cells) of LAAO^WT^ during a time course. Confocal live images show for each sample: LC3 EGFP fluorescence (top row), LC3 mCherry fluorescence (middle row) and the merged image (bottom row). Nuclei were stained with low concentration of Hoechst 33342 (blue), and images captured every 30 min. The images complement the data shown in Figure 4A-C and were collected in parallel with the same settings. Scale bar: 10 µm. N=3.

**Fig. S4.**
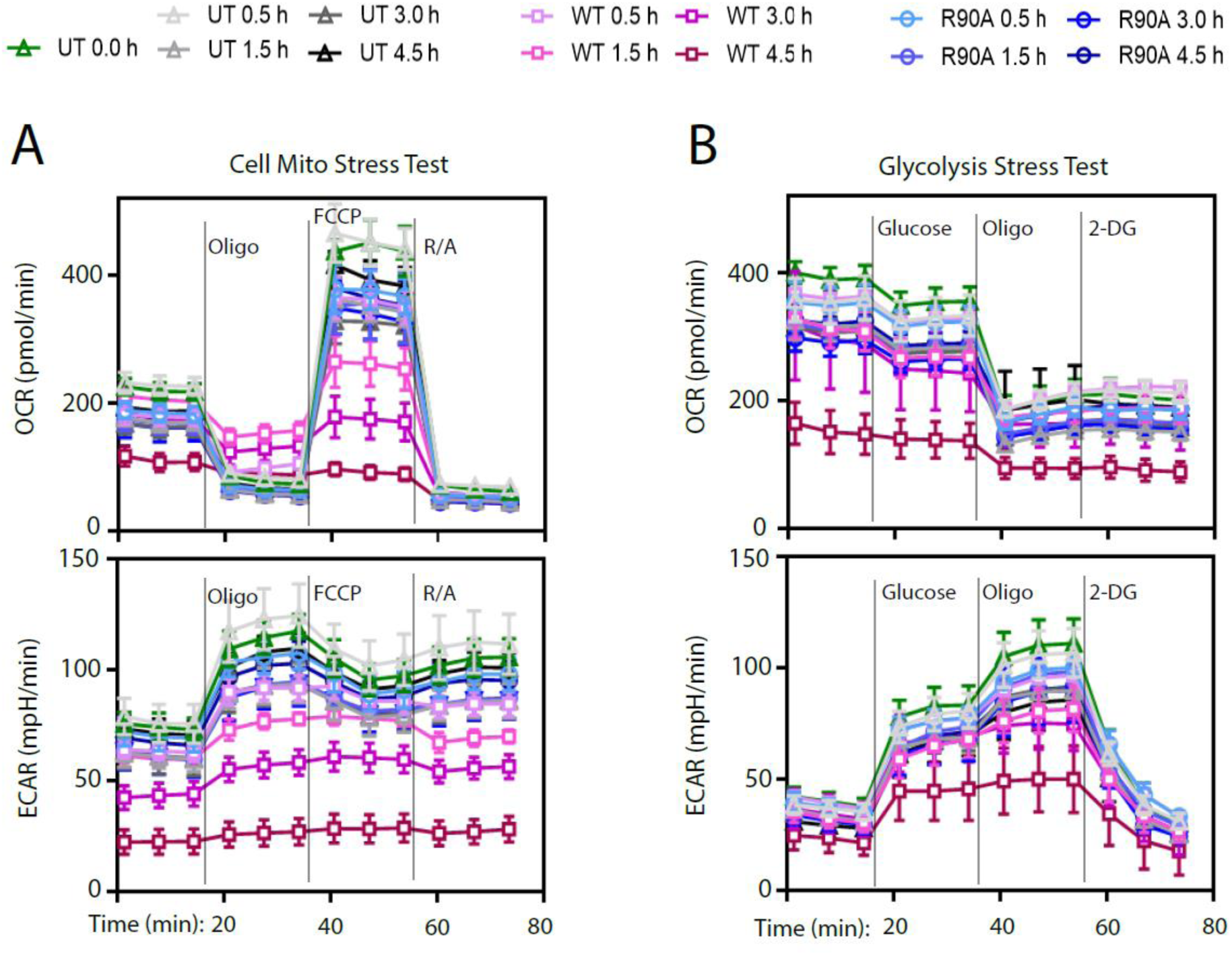
Profiles of oxygen consumption rate (OCR) and extracellular acidification rate (ECAR) following incubation with *B. atrox* recombinant LAAO^WT^ or LAAO^R90A^. Keratinocyte were left untreated (UT) or incubated with 2 X EC_50_ (73 nM for SeaHorse experiments) LAAO^WT^ or LAAO^R90A^ for 0.5, 1.5, 3.0, and 4.5 h. Cells were then subjected to mitochondrial stress test (A) or Glycolysis stress test (B). The profiles shown complement data shown in Figure 6. **A**, Mitochondrial stress test when challenged by oligomycin (oligo, an ATP synthase inhibitor), carbonylcyanide-p-trifluoromethoxyphenyl hydrazine (FCCP, an electron transport chain uncoupler) or rotenone/antimycin A (R/A, inhibitors of electron transport chain complex I and III). The addition of each chemical at a particular time is indicated as vertical grey lines. **B**, Glycolysis stress test was performed in the presence of glucose (fuel for glycolysis), oligomycin or 2-deoxyglucose (2DG, an inhibitor of glycolysis). N=3.

**Fig. S5.**
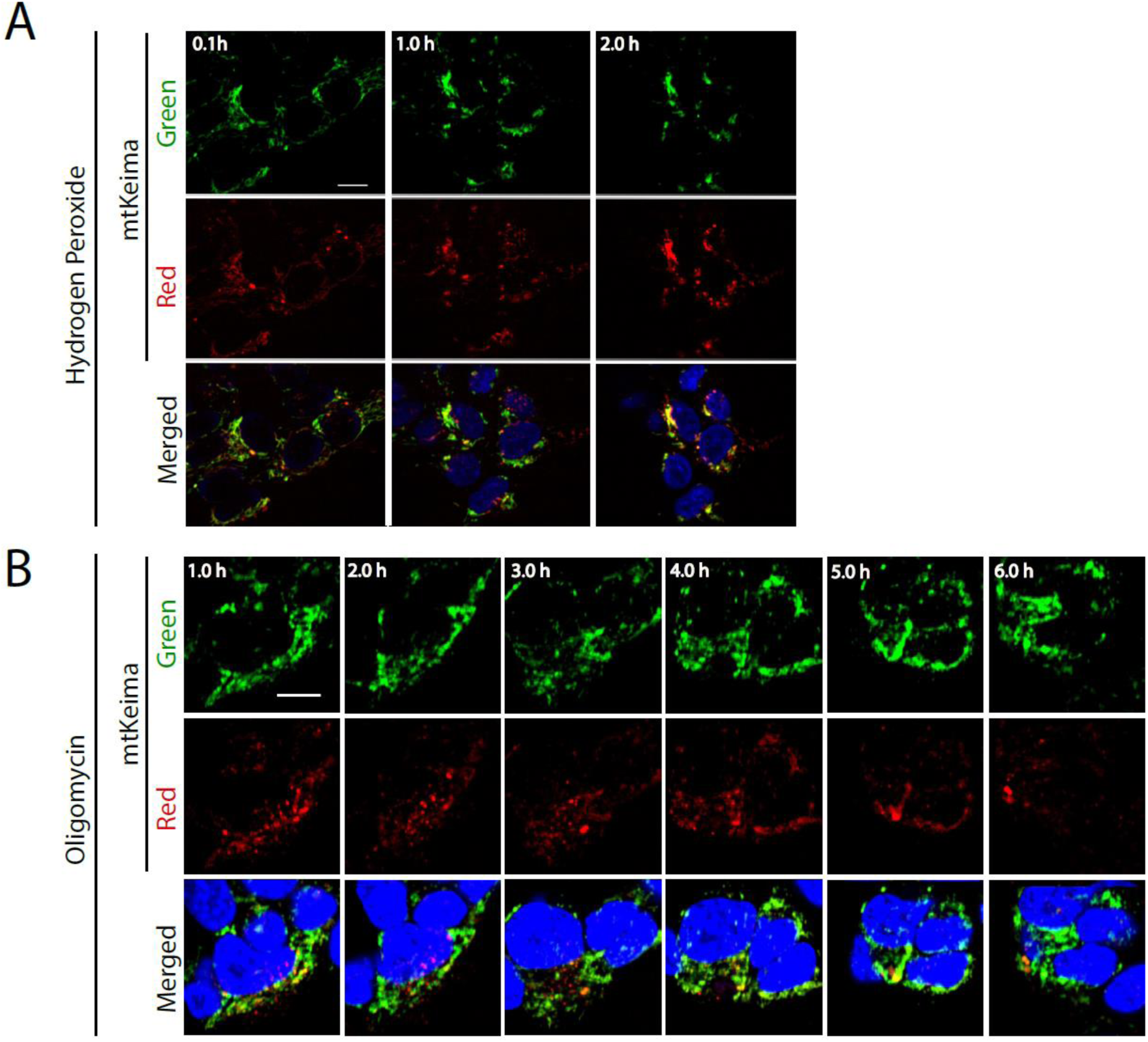
Mitophagy responses of SH-SY5Y cells stably expressing mtKeima to hydrogen peroxide (A) or oligomycin (B). Confocal live imaging of cells shows for each treatment: cytoplasmic mtKeima (green, top row), lysosomal mtKeima (red, middle row) and nucleus (blue, bottom row). The images complement the data shown in Figure 7C and were collected in parallel with the same settings. Scale bar: 10 µm. N=3.

**Table S1.**
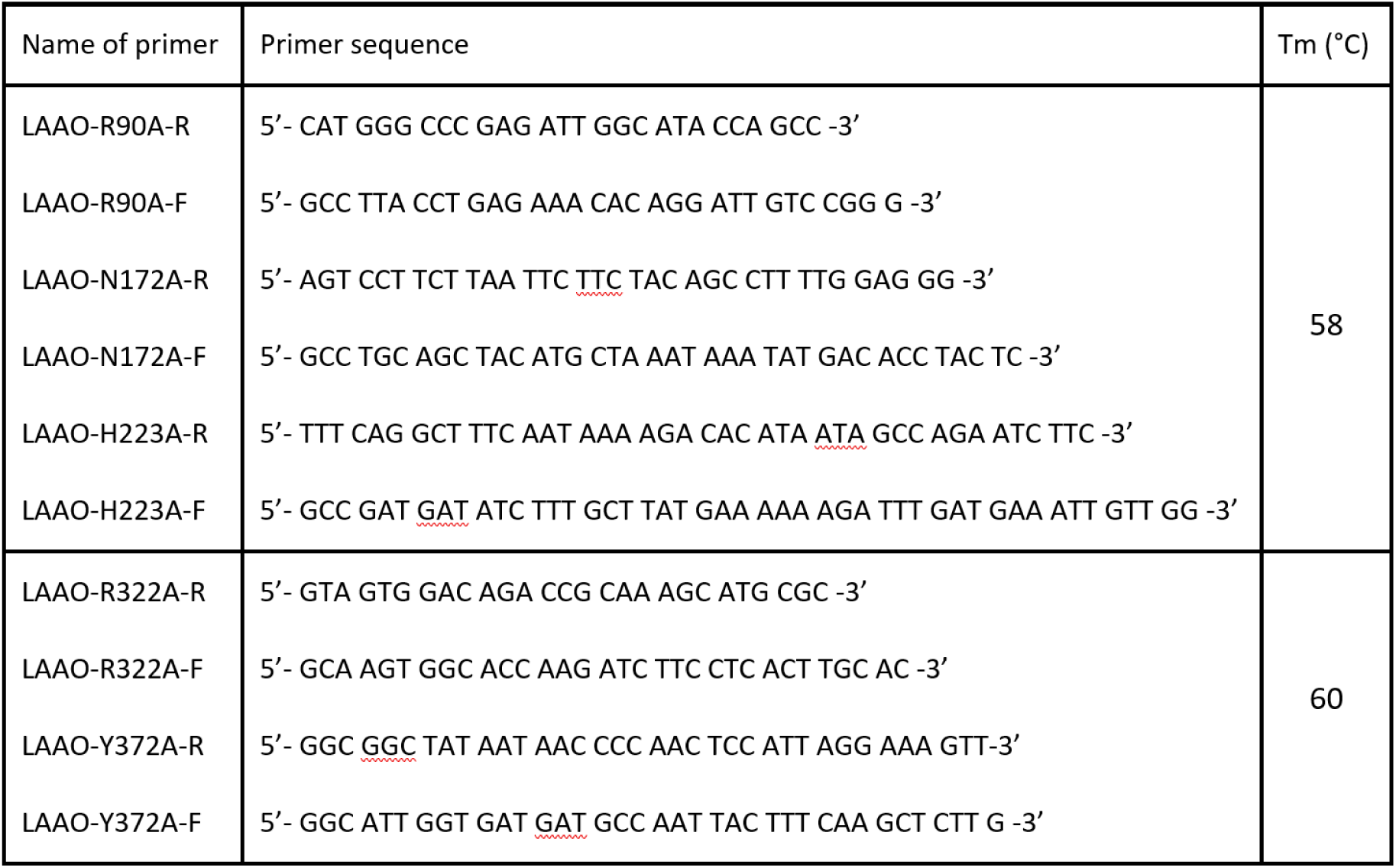
Primer sequences used for cloning.

